# Human P2X4 receptor gating is modulated by a stable cytoplasmic cap and a unique allosteric pocket

**DOI:** 10.1101/2024.07.25.605151

**Authors:** Haoyuan Shi, Ismayn A. Ditter, Adam C. Oken, Steven E. Mansoor

## Abstract

P2X receptors (P2XRs) are a family of ATP-gated ion channels comprising homomeric and heteromeric trimers of seven subunits (P2X_1_ – P2X_7_) that confer different rates of desensitization. The helical recoil model of P2XR desensitization proposes the stability of the cytoplasmic cap sets the rate of desensitization, but timing of its formation is unclear for slow-desensitizing P2XRs. We report cryo-EM structures of full-length, wild-type human P2X_4_ receptor in apo, antagonist-bound, and desensitized states. Because the apo and antagonist-bound structures of this slow-desensitizing P2XR include an intact cytoplasmic cap while the desensitized state structure does not, the cytoplasmic cap forms before agonist binding. Furthermore, structural and functional data suggests the cytoplasmic cap is stabilized by lipids to slow desensitization and that P2X_4_ is further modified by glycosylation and palmitoylation. Finally, our antagonist-bound inhibited state structure reveals features specific to the allosteric ligand-binding pocket in human receptors that empower the development of small-molecule modulators.

## Introduction

P2XRs are a family of trimeric, ATP-gated, non-selective cation channels that are broadly distributed across cells of the nervous, cardiovascular and immune systems (*1*–*7*). The continually emerging role of extracellular ATP as a signaling molecule in physiological and pathophysiological states has implicated P2XRs in vital processes such as platelet activation, synaptic transmission and inflammation, making P2XRs important pharmacological targets for a broad range of diseases (*3*, *4*, *8*–*12*). The functional diversity of the seven P2XR subunits results in receptors with varied sensitivities to ATP, unique susceptibilities to small-molecule agonists and antagonists, and different rates of desensitization following activation (*5*, *6*, *10*, *13*–*15*). Understanding the foundations of these subtype-specific properties is important for the development of small-molecule modulators targeted to each P2XR subtype.

Kinetics of channel gating by ATP is specific for each P2XR, allowing categorization into three broad groups: P2X_1_ and P2X_3_ receptors display fast rates of desensitization (millisecond timescales), P2X_2_, P2X_4_ and P2X_5_ receptors desensitize slowly (seconds), and P2X_7_ receptors do not desensitize at all (*5*, *10*, *13*). Insights into the molecular mechanisms of gating have been gained from structural investigations of the fast-desensitizing human P2X_3_ (hP2X_3_) receptor, which resulted in the ‘helical recoil’ model that describes the conformational changes associated with P2XR desensitization (*16*, *17*). Crystal structures of hP2X_3_ revealed that cytoplasmic residues in the N- and C-termini interact to form a domain, termed the cytoplasmic cap, that was structured in the ATP-bound open state but disordered in both the apo closed and ATP-bound desensitized states (*16*). Thus, transient formation of the cytoplasmic cap was proposed to provide a structural scaffold in the ATP-bound activated conformation to stabilize the open pore; its subsequent disassembly drives the transition to an ATP-bound desensitized conformation. In this framework, the stability of the cytoplasmic cap sets the rate and extent of receptor desensitization: fast-desensitizing P2XRs (P2X_1_ and P2X_3_) have a less stable cytoplasmic cap and slow-desensitizing subtypes (P2X_2_, P2X_4_, and P2X_5_) have a more stable cytoplasmic cap. Subsequent studies of the non-desensitizing rat P2X_7_ (rP2X_7_) receptor revealed the presence of the cytoplasmic cap in the apo closed state structure, permanently stabilized by a series of P2X_7_-specific palmitoylated residues that are crucial for preventing desensitization (*18*), consistent with the notion that the stability of the cytoplasmic cap sets the rate and extent of desensitization. While there is structural evidence explaining the gating kinetics for fast-desensitizing (hP2X_3_) and non-desensitizing (rP2X_7_) P2XRs, the structural basis for slow-desensitizing P2XRs remains poorly understood (*16*, *18*).

The P2X_4_ receptor subtype, which undergoes slow desensitization, is closely related to P2X_7_, both structurally and functionally (*19*–*21*). Initial insights into the architecture of P2X_4_ receptors were provided by apo closed and ATP-bound open state structures of zebrafish P2X_4_ (zfP2X_4_) receptor truncated at both the N- and C-termini (*22*, *23*). More recently, structures of zfP2X_4_ bound to allosteric inhibitors, BAY-1797 and BX430, were resolved using similarly truncated constructs, revealing allosteric ligand-binding sites in analogous locations to those found in P2X_7_ (*24*–*26*). Truncations to zfP2X_4_ were important for protein overexpression and stability but led to uncertainty about the structure of its cytoplasmic domain. As a direct result, it is unknown whether the cytoplasmic cap is present in the apo closed and ATP-bound open states of P2X_4_ (as for P2X_7_) or if it assembles upon transition from the apo closed state to the ATP-bound open state (as for P2X_3_). Thus, it is unclear exactly how the helical recoil model applies to slow-desensitizing P2XR subtypes and whether there are any additional structural determinants that dictate slow desensitization.

Here we present three high-resolution cryo-EM structures of the full-length wild-type human P2X_4_ (hP2X_4_) receptor in the apo closed state, ATP-bound desensitized state, and allosterically inhibited state bound to BAY-1797 (N-[4-(3-Chlorophenoxy)-3-sulfamoylphenyl]-2-phenylacetamide) (*27*). In contrast to previous P2X_4_ structures, the cytoplasmic cap is resolved in the apo closed state where it is stabilized by putative lipid-binding sites, revealing a distinguishing feature for slow-desensitizing P2XR subtypes. The ATP-bound desensitized state structure lacks an assembled cytoplasmic cap, and thus provides information on the overall architectural changes following receptor desensitization, notably the changes to the pore compared to the apo closed state. The BAY-1797-bound inhibited state of hP2X_4_ preserves structural elements found in the apo closed state, including the pore architecture and the cytoplasmic cap. Further, the BAY-1797-bound inhibited structure defines human-specific features of the P2X_4_ allosteric ligand-binding pocket. Together, these data provide critical insights into the mechanisms underlying the distinct desensitization profiles of different P2XR subtypes and present an opportunity to target the human ortholog to develop small molecules for therapeutic use.

## Results

### Overall architecture of the apo closed state of hP2X_4_ receptor

To improve our understanding of homomeric P2X_4_ receptors, we obtained cryo-EM structures of the full-length wild-type hP2X_4_ receptor in multiple conformations (Fig. S1,S2, Table S1). As expected, the architecture of hP2X_4_ resembles other P2XRs, with domains in each protomer named according to the anatomy of a breaching dolphin, as previously described (Fig. S3) (*16*, *18*, *22*). Each protomer of hP2X_4_ is comprised of a large, hydrophilic extracellular domain (ECD), two transmembrane (TM) spanning alpha helices (the outer TM1 and pore-lining TM2), and intracellular N- and C-termini (Fig. S3). Unexpectedly, however, the apo closed state structure of hP2X_4_ has a cytoplasmic cap, redefining our understanding of the P2XR gating cycle for at least this P2X subtype (Fig. 1, 2, S3, S4A,B).

**Fig. 1.**
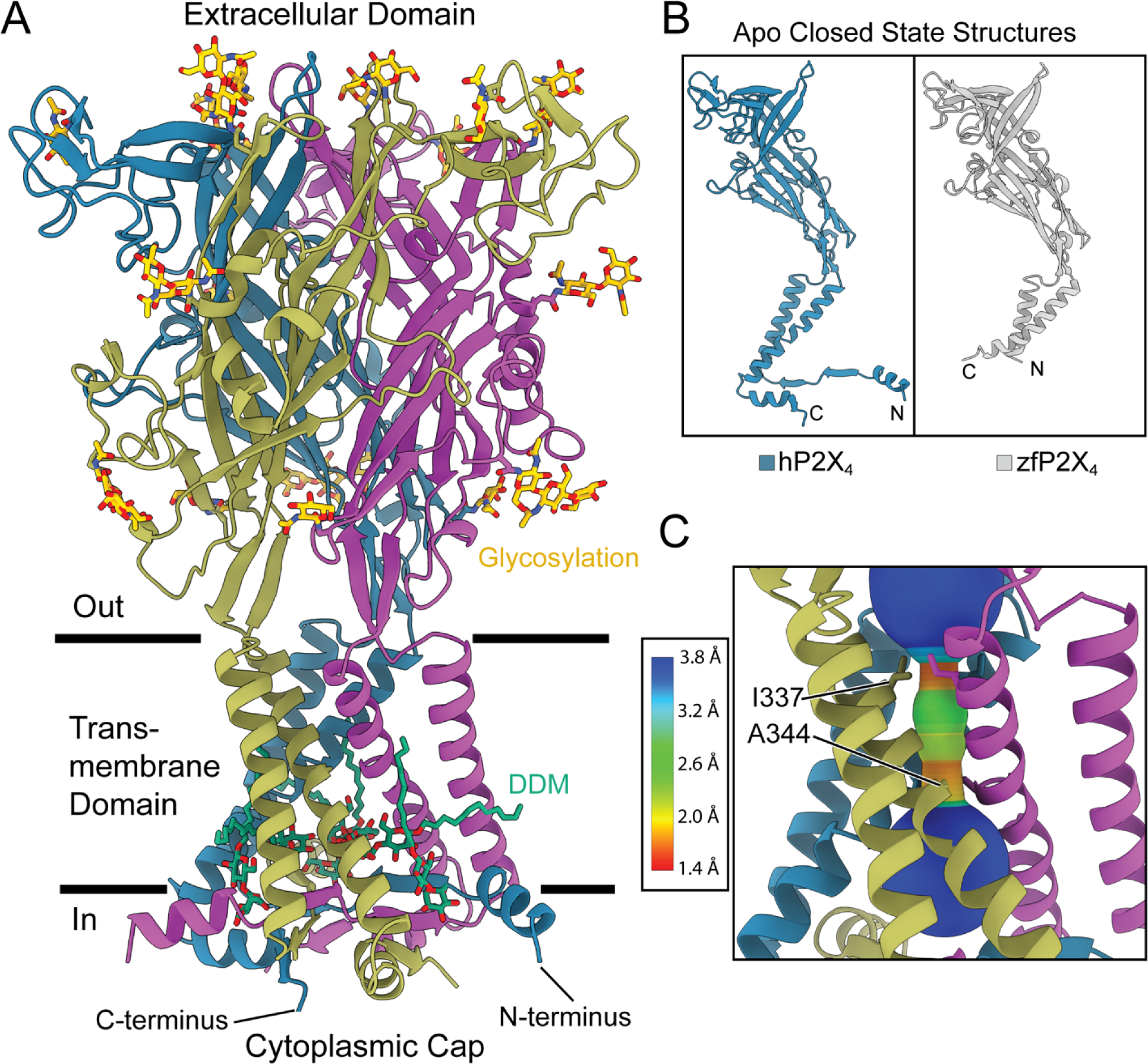
Overall architecture and pore structure of hP2X_4_ in the apo closed state. **(A)** Ribbon representation of the full-length, wild-type human P2X_4_ receptor in the apo closed state, highlighting the presence of the cytoplasmic cap, multiple glycosylation sites in the extracellular domain, and putative lipid-binding sites in the cytoplasmic domain. Each protomer of the receptor is a different color: blue, purple, and dark yellow. Glycosylations are shown in yellow. Dodecyl maltoside (DDM) molecules are shown in green. **(B)** Ribbon representation of a single protomer of hP2X_4_ in the apo closed state compared to the previously published structure of zfP2X_4_ in the apo closed state, highlighting how the new human structure is significantly more complete with TM domains that span the lipid bilayer and contains the cytoplasmic cap. **(C)** The ion permeation pathway of hP2X_4_ in the apo closed state, revealing that the constriction sites occur at two distinct residues on TM2: an initial gate formed by I337 from each protomer (1.0 Å pore radius) and then a tighter secondary gate formed by A344 from each protomer (0.4 Å pore radius). For the pore size plot, different colors represent the pore radius, as generated by the program MOLEonline (*45*).

**Fig. 2.**
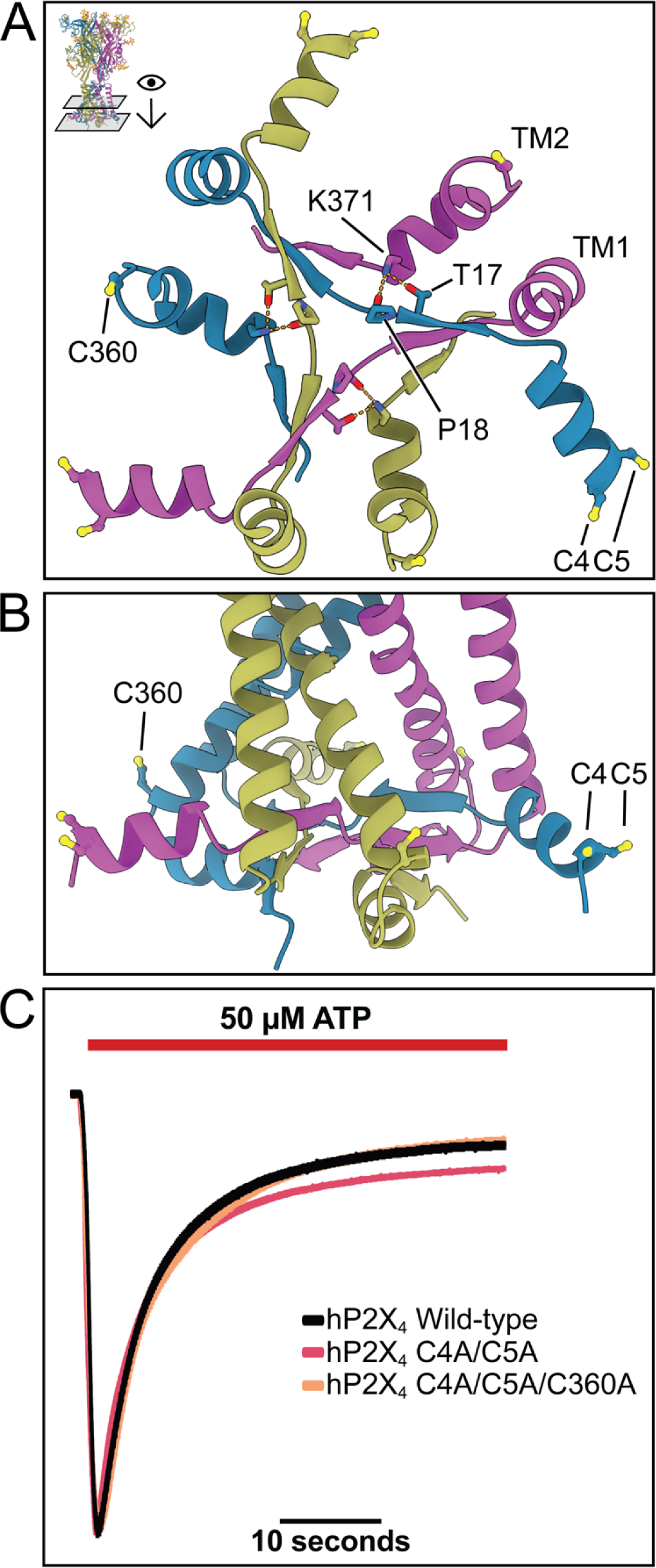
The cytoplasmic cap of hP2X_4_ is palmitoylated, but these post-translational modifications do not modulate the rate of receptor desensitization. **(A)** Top-down view of the cytoplasmic cap of hP2X_4_, with residue interactions important for the cap structure shown as yellow dashed lines. The location and orientation of two cysteine residues in the N-terminus (C4 and C5) and one cysteine residue in the C terminus (C360) are labeled. Mass spectrometry analysis confirmed that residues C4 and C5 can be palmitoylated. Peptide fragments containing C360 could not be identified in the mass spectrometry studies, but a similarly located residue in P2X_7_ receptor is palmitoylated. **(B)** Side-view of the TM domain showing how the cytoplasmic cap is composed of domain-swapped β-strands from each protomer. Residues C4, C5, and C360 each face toward the lipid bilayer. **(C)** Preventing palmitoylation by mutation of residues in hP2X_4_ confirmed to be palmitoylated (C4, C5) or have the potential to be palmitoylated (C360) results in mutant hP2X_4_ receptors (C4A/C5A and C4A/C5A/C360A) that, when expressed in *Xenopus* oocytes, possess desensitization kinetics in response to ATP that are indistinguishable from wild-type hP2X_4_ receptor. These TEVC findings suggest that palmitoylation of residues in hP2X_4_ do not modulate the rate of receptor desensitization.

The cryo-EM structure of hP2X_4_ in the apo closed state at 2.3 Å resolution, defined by an empty orthosteric ligand-binding pocket and a closed channel pore, significantly improves upon previous P2X_4_ structures, revealing the structure of the cytoplasmic domain and several subtype-specific features (Fig. 1A,B, Fig. 2A,B) (*22*–*24*). For example, putative lipid-binding sites and significant N-linked glycosylation are readily visualized. In addition, numerous water molecules can be placed throughout the receptor, especially in ligand-binding sites, facilitating small-molecule ligand development (*28*, *29*). Finally, we provide the first structure of the human P2X_4_ receptor, enabling comparisons between previously characterized orthologs as well as empowering structure-based drug design for the most pharmacologically relevant species. The structure of the ECD is highly conserved between human and zebrafish P2X_4_ orthologs (*23*, *24*). For instance, alignment of the ECD between the apo closed state structures of hP2X_4_ (residues 50-331) and zfP2X_4_ (PDB: 4DW0, residues 53-334) yielded a backbone RMSD of only 0.780 Å (*23*). However, we observed significant differences within the remaining domains, identifying novel structural features.

As a result of using the full-length hP2X_4_ construct, the entire conductance pathway is visualized for the first time. The conductance pathway of hP2X_4_ is blocked at two locations: an initial constriction gate formed by I337 from each protomer (1.0 Å pore radius), followed below by a tighter constriction gate formed by A344 from each protomer (0.4 Å pore radius) (Fig. 1C, Fig. S4C). These gate residues correspond to L340 (0.7 Å pore radius) and A347 (0.7 Å pore radius) in zfP2X_4_ (*23*) (Fig. S4D). The channel is defined as closed since the pore radius is too narrow to pass dehydrated Na^+^ ions (*30*). Below the gate, the structure of hP2X_4_ diverges from zfP2X_4_ (Fig. 1B, S4). In zfP2X_4_, each TM2 helix differs in pitch from hP2X_4_ at the cytoplasmic end by a relatively small angle (10.7°) (Fig. S4D). In contrast, each TM1 helix is dramatically bent in zfP2X_4_, with the last four turns of the helix pointing away from the receptor pore, altering the pitch by 38.3° compared to TM1 in hP2X_4_ (Fig. S4D). The disparate orientations of the TM domains are likely explained by the presence of the cytoplasmic cap in the full-length structure of hP2X_4_, where it constrains the TM domain and thus sets its orientation and pitch (Fig. 1B, Fig. S4).

### The cytoplasmic cap is assembled in the apo closed state structure of hP2X_4_

Structural studies of wild-type hP2X_4_ allow visualization and modeling of nearly the entire receptor: the apo closed state model begins at residue G3 and ends at residue E376 (374 of 388 total residues are present in the model) (Fig. 1A,B). As a result, we visualize the complete trans-membrane domain of hP2X_4_, presenting a model that fully spans the membrane bilayer (Fig. 1A). Furthermore, we resolve residues within the cytoplasmic domain of hP2X_4_ that compose the cytoplasmic cap, a structural feature now seen in hP2X_4_ for the first time (Fig. 1A,B, 2A). The cytoplasmic cap (first described in the ATP-bound open state structure of hP2X_3_) is formed by residues from both the N- and C-termini of the receptor, where the three protomers intertwine to form three β-sheets that run parallel to the membrane (*16*). Each β-sheet contains a β-strand from each monomer, linked by 9 sets of hydrogen bonds formed by the protein backbone (Fig. S5A). In addition to backbone interactions, specific sidechains in the cytoplasmic cap also form hydrogen bonds (Fig. 2A). Residue P18 alters the peptide backbone geometry of the β-strand and orients its carbonyl oxygen to form a hydrogen bond with K371 (2.8 Å) (Fig. 2A). Additionally, K371 forms a hydrogen bond with T17 (2.7 Å) (Fig. 2A). These sets of hydrogen bonds, both from the backbone interactions within the β-strands and between sidechains, likely form the structural basis necessary for stability of the cytoplasmic cap.

The cytoplasmic cap in the apo closed state structure of hP2X_4_ bears similarities to the cytoplasmic cap observed in the ATP-bound open state structure of hP2X_3_ (*16*). The ATP-bound open state structure of hP2X_3_ was obtained using a P2X_2_/P2X_3_ chimeric construct containing three key mutations in the cytoplasmic cap: T13P, S15V, and V16I. Mutation of these three key residues in the N-terminus of hP2X_3_ receptor to the residues that exist in P2X_2_ receptor was demonstrated to slow desensitization of this normally fast-desensitizing P2XR subtype by providing stabilizing hydrophobic interactions in the cytoplasmic cap (*16*, *31*). The three aforementioned residues correspond to P18, I20, and V21 in hP2X_4_, respectively, where they are located in similar positions in both structures. Furthermore, the previously discussed residues K371 and T17 in hP2X_4_ (corresponding to K357 and T12 in hP2X_3_, respectively), are conserved in both orthologs. When aligning the cytoplasmic region of the chimeric hP2X_3_ in the ATP-bound open state structure (PDB: 5SVK) to hP2X_4_ in the apo closed state structure, each of these discussed residues overlap entirely, suggesting that they play a similar role in the hP2X_4_ apo closed state conformation to stabilize this domain of the receptor (Fig. S5B,C) (*16*).

### The cytoplasmic cap is palmitoylated

While analyzing the apo closed state structure of hP2X_4_, we observed three cysteine residues per protomer (C4, C5, and C360) pointing up toward the lipid bilayer (Fig. 2A,B), in positions strikingly similar to cysteine residues previously documented to be palmitoylated in rP2X_7_ (*18*). Although definitive density for palmitoyl moieties are not present in our cryo-EM reconstructions of hP2X_4_, mass spectrometry (MS) analysis confirms that at least two palmitoylations sites (C4 and C5) are present in hP2X_4_ - these two cysteine residues are located on an α-helix in the cytoplasmic cap at the N-terminus of the receptor and point upward and outward toward where the plasma membrane would likely be located (Fig. 2A,B). MS further confirms that three combinations of post-translational modifications are present: single palmitoylation at cysteine 4, single palmitoylation at cysteine 5, and double palmitoylation at both C4 and C5 (Fig. S6). Unmodified peptides were also detected suggesting that palmitoylation is not stoichiometric. In the case of singly palmitoylated peptides, there appears to be no preference for either cysteine, as intensities from MS/MS spectra were similar in intensity (Fig. S6).

To evaluate the role of palmitoylation on receptor function and gating, two hP2X_4_ mutants (C4A/C5A and C4A/C5A/C360A) were generated to prevent palmitoylation and tested by two-electrode voltage clamp (TEVC) experiments on *Xenopus* oocytes (Fig. 2C). Of note, we were unable to detect the peptide fragment containing residue C360 during MS analysis and thus, we could not determine whether it is palmitoylated. However, based on its location and homology to a palmitoylation site in rP2X_7_, we elected to include it in the functional analysis (*18*). When activated by extracellular ATP, both mutants featured fast activation followed by slow desensitization, mirroring the kinetics of wild-type hP2X_4_ receptor (Fig. 2C). Normalization of the mutant and wild-type current traces reveals similarities in the activation and desensitization kinetics. Additionally, the current amplitudes were essentially equivalent between mutant and wild-type receptors, suggesting that expression and trafficking were not significantly affected by removing the palmitoyl groups. Although it is currently unclear why residues C4 and C5 in hP2X_4_ are palmitoylated, they do not appear to play a significant role in modulating receptor gating.

### The cytoplasmic cap contains putative lipid-binding sites

The apo closed state reconstruction of hP2X_4_ contains unexpected nonprotein densities within the cytoplasmic cap (Fig. S7A). The size and shape of the densities are consistent with molecules of dodecyl maltoside (DDM), the detergent used to solubilize and reconstitute the receptor (Fig. S7B,C). Two DDM molecules per protomer (six total in the receptor) are present in the model (Fig. 3A). The first DDM molecule is positioned vertically at the monomer interface, where the maltoside group makes interactions with residues on the receptor termini and the dodecyl chain extends upwards alongside the TMs, parallel to the axis of symmetry (Fig. 3A,B). The sugar head group from this DDM molecule forms hydrogen bonds to the sidechains of Y359, Y372, R33, R368, and E14 (3.4 Å, 3.0 Å, 3.2 Å, 3.0 Å, and 2.9 Å, respectively) (Fig. 3B). Additionally, the dodecyl moiety forms hydrophobic interactions, extending deep into the detergent micelle, with residues I339 and I346 on TM2 (3.3 Å and 3.2 Å, respectively) as well as residue V43 on TM1 (3.4 Å). A second DDM molecule is placed perpendicular to the axis of symmetry and adjacent to the first DDM molecule (Fig. 3A,C). The sugar head groups from this DDM molecule form hydrogen bonds with a water molecule and the sidechains of D16 and D354 (3.0 Å, 3.0 Å, and 3.0 Å, respectively) (Fig. 3C). The dodecyl moiety forms hydrophobic interactions with L37 (3.9 Å) on TM1. Hydrogen bonds between the DDM molecules are also observed (Fig. 3A).

**Fig. 3.**
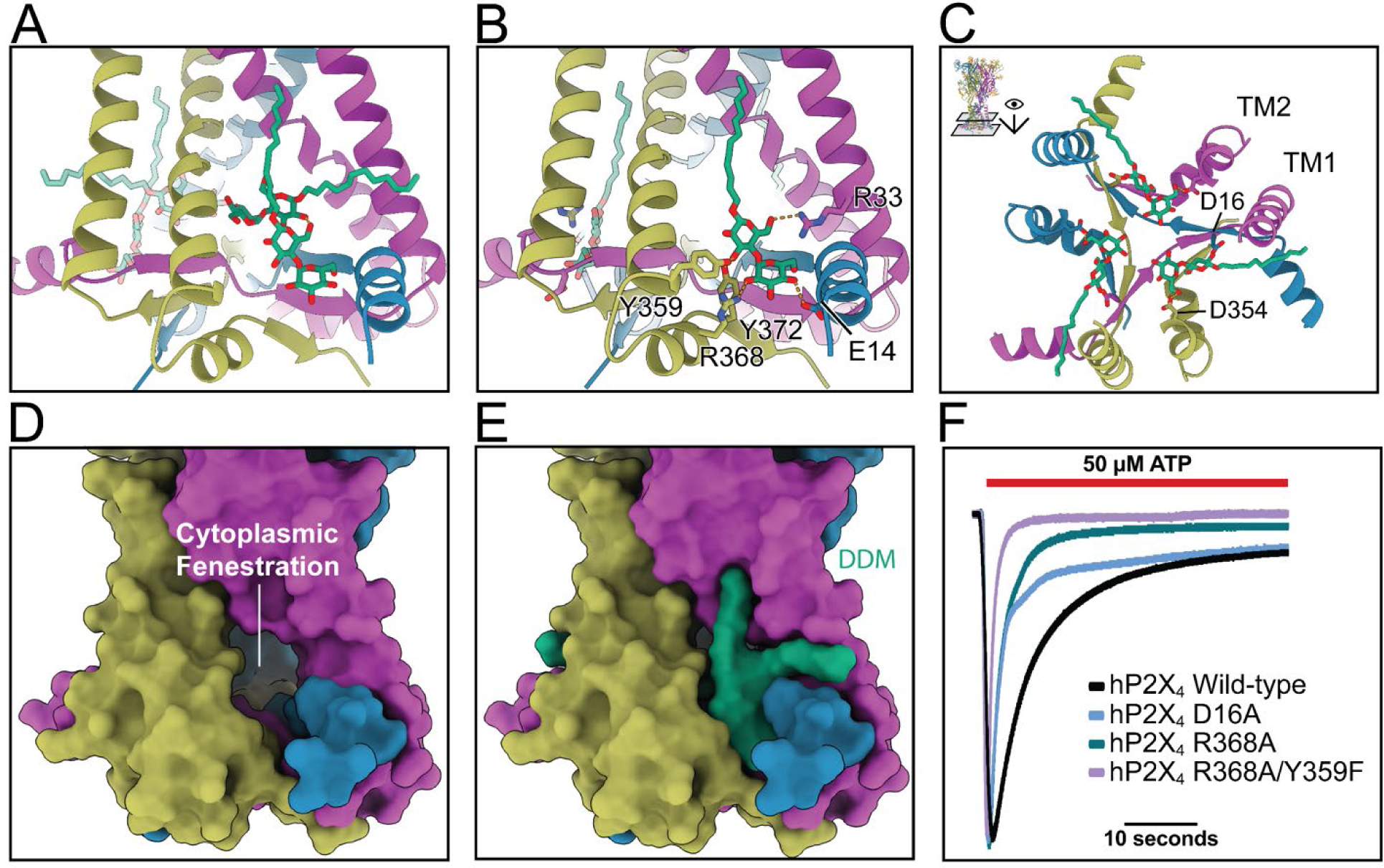
The cytoplasmic cap of hP2X_4_ has putative lipid-binding sites that modulate the rate of receptor desensitization. **(A)** View of the putative lipid-binding site above the cytoplasmic cap of hP2X_4_, with two DDM detergent molecules bound per protomer (shown in green). Interactions between the DDM molecules consist of hydrogen bonding between the O6 on the first sugar of the vertically positioned DDM and the O2 and O6 of the first sugar on the horizontally positioned DDM (3.0 Å and 3.2 Å, respectively). **(B)** View of the vertical DDM molecule highlighting interactions it makes to residues that compose the cytoplasmic cap and the cytoplasmic fenestration. On the first glucose, O3 hydrogen bonds to the sidechains Y359 and Y372 of one protomer and O6 hydrogen bonds to the sidechain of R33 of another protomer (H-bond distances of 3.4 Å, 3.0 Å, and 3.2 Å, respectively). The second glucose also forms hydrogen bonds, where O2 interacts with the sidechain of R368 and O4 with the sidechain of E14 (H-bonds distances of 3.0 Å and 2.9 Å, respectively) **(C)** Top-down view of the cytoplasmic cap highlighting the horizontal DDM molecule and the interactions it makes to the receptor. On the first glucose, O3 forms hydrogen bonds with the sidechain of D16 (3.0 Å). The O6 of the second sugar forms hydrogen bonds with the sidechain of D354 (3.0 Å) **(D)** Space filling view of the cytoplasmic fenestration in hP2X_4_ receptor with DDM molecules hidden. Molecular dynamics simulations in hP2X_3_ receptor suggest the cytoplasmic fenestrations are water-filled rivulets juxtaposed between the protein and lipid membrane that function as pathways for ion egress into the cytoplasm. **(E)** Space filling view of the cytoplasmic fenestration in hP2X_4_ receptor showing how the two DDM molecules bind in the cytoplasmic cap at the interface between transmembrane helices of different monomers. **(F)** Mutation of residues in putative lipid-binding sites (D16A, R368A, and R368A/Y359F) within the cytoplasmic domain of hP2X_4_ results in significantly increased rates of desensitization compared to wild-type hP2X_4_ receptor in TEVC experiments. The electrophysiology data suggest that the detergent-binding sites discovered in the structure of hP2X_4_ receptor likely represent lipid-binding sites *in vivo* that modulate hP2X_4_ function by affecting the stability of the cytoplasmic cap.

Both DDM molecules bind to residues on the cytoplasmic cap as well as residues at the interface between transmembrane helices of different monomers, at what has been referred to in P2X receptors as the cytoplasmic fenestrations (Fig. 3D, E) (*16*). Molecular dynamics studies in hP2X_3_ suggest that the cytoplasmic fenestrations are a lipid-lined path for water and ions to egress from the channel into the cell (*16*). Since P2X_4_ receptor activity has previously been shown to be regulated by phosphoinositides through direct interactions in the C-terminal domain, we wanted to explore the possibility that these binding pockets occupied by DDM in our structures of hP2X_4_ represent lipid-binding sites *in vivo* (*32*).

The role of these putative lipid-binding sites on receptor function and gating was elucidated by assessing the gating properties of hP2X_4_ mutants (D16A, R33A, R368A, and R368A/Y359F) predicted to have a reduced ability to bind putative lipids, based on residue interactions between DDM and the receptor. In TEVC experiments, altering residues within the putative lipid-binding sites either abrogated receptor function or significantly impacted gating by increasing the rate of receptor desensitization (Fig. 3F). For receptors containing the R33A mutation, we observed negligible ATP-induced current (data not shown). On the other hand, receptors containing the mutation D16A, R368A, or the double mutation R368A/Y359F possessed altered gating but with similar ATP-induced current intensity to the wild-type receptor. The wild-type hP2X_4_ receptor typically has a desensitization lifetime (τ) of ∼6.5 seconds (s), which describes the time it takes for the current to return to 63% of baseline during continuous application of extracellular ATP (Fig. 2C,3F). In contrast, the D16A, R368A, and R368A/Y359F mutants featured a fast activation and a fast desensitization that more resembled the gating kinetics of the fast-desensitizing P2X_3_ receptor (Fig. 3F). The single mutations D16A and R368A increased the rate of desensitization (τ = ∼3 s and ∼2.5 s, respectively), and the R368A/Y359F double mutant exhibited an even faster rate of receptor desensitization (τ = ∼1 s) (Fig. 3F). In addition, the R368A single mutant nearly completely and the R368A/Y359F double mutant completely desensitizes back to baseline current (prior to the addition of ATP), as opposed to wild-type hP2X_4_ which always exhibits a residual inward current even after very prolonged exposure to ATP (Fig. 3F). These functional data suggest that residues within the putative lipid-binding sites affect the stability of the cytoplasmic cap and thus modulate hP2X_4_ gating.

### The ATP-bound desensitized state structure of hP2X_4_ lacks the cytoplasmic cap

We report the cryo-EM structure of hP2X_4_ in the ATP-bound desensitized state at a resolution of 2.4 Å (Fig. 4A-D, Fig. S1, S2, Table S1). The structure of this conformational state was acquired from the same cryo-EM grid used to obtain the aforementioned apo closed state structure. As such, the purified protein sample was not incubated with any additional ATP during protein purification or prior to vitrification. Presumably, endogenous ATP binds to the receptor following cell lysis and remains bound throughout the purification process and structural analysis. This finding was observed previously for the hP2X_3_ receptor (*16*). During 3D reconstruction of the cryo-EM data, we were able to separate the apo closed state particles from the ATP-bound desensitized state particles based on the presence of the cytoplasmic cap, which is absent in the desensitized state (Fig. S1).

**Fig. 4.**
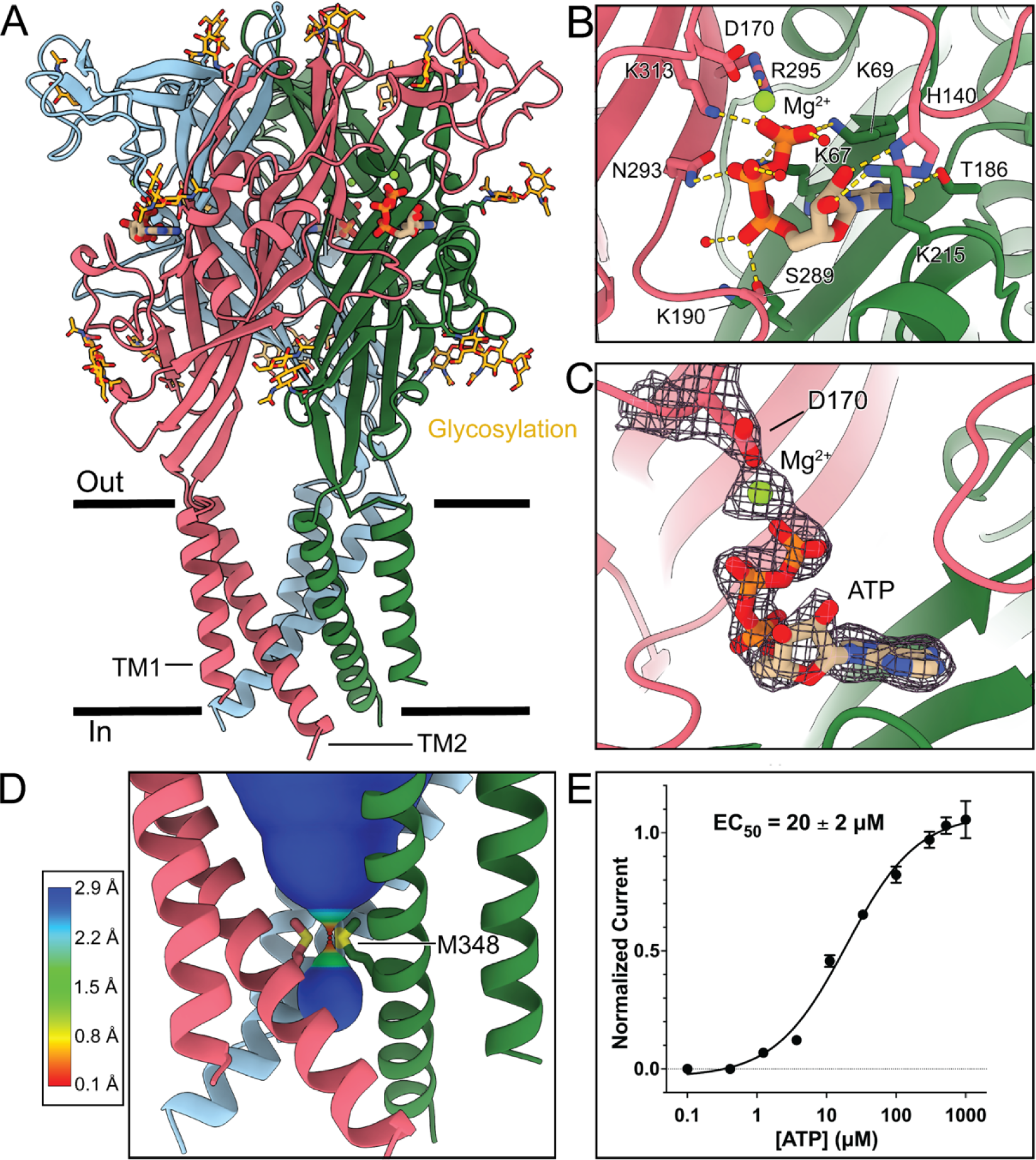
The cytoplasmic cap is disassembled in the ATP-bound desensitized state structure of hP2X4. **(A)** Ribbon representation of full-length, wild-type hP2X_4_ receptor in the ATP-bound, desensitized state, where the cytoplasmic cap is disassembled and not visible. Each protomer of the receptor is a different color: salmon, light blue, and green. Glycosylations are shown in yellow. ATP molecules are colored by atom: carbon atoms shown in tan, oxygen atoms in red, nitrogen atoms in blue, and phosphorus atoms in orange. Mg^2+^ ions are shown in lime green. **(B)** Close up view of the orthosteric ATP-binding site. Hydrogen bonds between ATP and residues are shown in yellow dotted lines. Of note, an Mg^2+^ ion was resolved within the orthosteric pocket. **(C)** Alternate view of the orthosteric pocket, where densities for D170, Mg^2+^ ion, and ATP are shown. **(D)** The ion permeation pathway of hP2X_4_ in the ATP-bound desensitized state, revealing that the constriction (gate) occurs at M348 on TM2 from each protomer (0.13 Å pore radius). Compared to the gate in the apo closed state shown in Fig. 1, the gate in the ATP-bound desensitized state is deeper into the membrane bilayer. For the pore size plot, different colors represent the pore radius, as generated by the program MOLEonline (45). **(E)** The apparent affinity (EC_50_) of ATP for wild-type hP2X_4_ is measured to be 20 ± 2 µM from two electrode voltage clamp (TEVC) experiments in Xenopus oocytes. The Y-axis describes currents normalized to a preceding current evoked by repeated applications of 300 µM ATP. The reported EC_50_ and error bars represent the mean and standard deviation across triplicate experiments, respectively.

We determined the apparent affinity of ATP (EC_50_) to hP2X_4_ to be 20 ± 2 µM by TEVC, consistent with previous work (Fig. 4E) (*6*). With ATP bound in the orthosteric pocket and a uniquely closed pore, the structure is defined as desensitized. The orthosteric ATP-binding site of hP2X_4_ is similar to other P2XRs, where seven conserved residues in the binding pocket form interactions with ATP: K67, K69, K190, K313, T186, N293, and R295 (Fig. 4B). Three additional residues, K190, S289, and H140, form subtype-specific hydrogen bonding interactions with the ATP (3.2 Å, 3.0 Å and 3.1 Å, respectively) (Fig. 4C). Specifically, the side chains of H140 and K215 form hydrogen bonds with the 2’ and 3’ hydroxyls on the ribose ring, respectively, while the side chain hydroxyl of S289 forms a hydrogen bond with an oxygen on the α-phosphate of ATP. Further, three water molecules within the orthosteric pocket were observed to form hydrogen bonds with the α-, β-, or γ-phosphates of the ATP (2.6 Å, 2.9 Å, and 3.1 Å, respectively). The quality of the reconstruction allowed the modeling of a Mg^2+^ ion coordinated by the γ-phosphate of ATP and the sidechain of D170 (Fig. 4C). Mg^2+^ is known to play a role in P2X_4_ function, where recent molecular dynamics studies suggested the presence of Mg^2+^ in the orthosteric pocket stabilizes the ATP-bound open state of zfP2X_4_ (*33*).

When comparing the overall structure of hP2X_4_ in the ATP-bound desensitized state to the apo closed state, we observed similarities in the ECD (residues 51-331, RMSD of 0.85 Å). Consistent with the helical recoil model of P2X receptor desensitization, the TM domains are displaced outwards and upwards, where the resulting conformational changes to TM2 alter the location of the constriction site in the pore (*16*, *17*). As a result, the new gate, now formed by M348 from each protomer (0.13 Å pore radius), is shifted deeper into the membrane bilayer compared to the gate in the apo closed state (Fig. 1C, Fig. 4D). The architecture of the TM domain in the ATP-bound desensitized state results from disassembly of the cytoplasmic cap, and consequently, the cytoplasmic cap is unresolved in the ATP-bound desensitized state structure (Fig. 4A). Residues 3-30 and 362-376 are missing in our cryo-EM reconstruction of hP2X_4_ in the ATP-bound desensitized state, suggesting flexibility or disorder. Similarly, no cytoplasmic cap was observed in the ATP-bound desensitized state structure of hP2X_3_, indicating the transition from open to desensitized conformational states likely follows parallel mechanisms in both hP2X_3_ and hP2X_4_.

### BAY-1797 binds to the allosteric site and preserves the cytoplasmic cap

We also report the cryo-EM structure of the allosteric antagonist BAY-1797 bound to hP2X_4_ at a resolution of 2.6 Å (Fig. 5, S1,S2,S8,S9, Table S1). BAY-1797 inhibits hP2X_4_ with a half-maximal inhibitory concentration (IC_50_) of 220 ± 30 nM as measured by TEVC experiments, consistent with previous electrophysiological data (Fig. 5B) (*24*). The overall receptor architecture in the antagonist-bound inhibited state resembles the apo closed state structure (RMSD of 0.5 Å) (Fig. S9), where many of the unique features are still present: the pore is closed with gates at residues I337 and A344, the cytoplasmic cap remains ordered and assembled, and the detergent molecules are bound in the cytoplasmic fenestrations (Fig. 5A). However, in comparison to the apo closed state structure, binding of BAY-1797 alters the local architecture of the allosteric site by expanding the pocket, conferring movements in the upper body domains to accommodate the ligand (Fig. S3, S9).

**Fig. 5.**
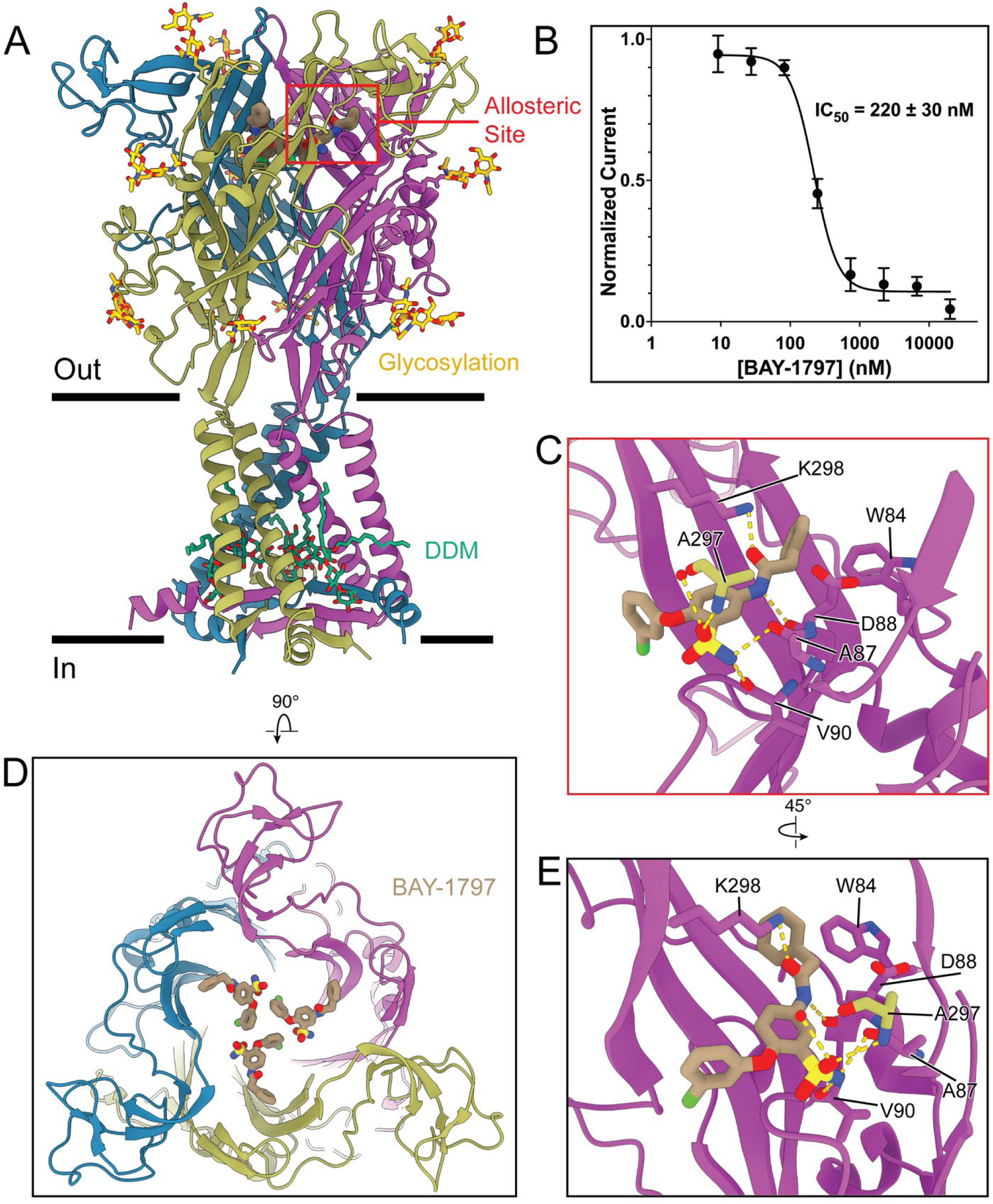
BAY-1797 binds to the allosteric binding site in hP2X4 and preserves the cytoplasmic cap. **(A)** Ribbon representation of BAY-1797 bound to full-length wild-type hP2X_4_ receptor. The allosteric binding pocket is highlighted with a red box. Relative to the apo closed state, the binding of BAY-1797 does not impact the TM architecture, the presence of the cytoplasmic cap, or the presence of putative lipid-binding sites in the cytoplasmic domain. Each protomer of the receptor is a different color: blue, purple, and dark yellow. Glycosylations are shown in yellow. BAY-1797 molecules are colored by atom: carbon atoms shown in tan, oxygen atoms in red, nitrogen atoms in blue, chlorine atoms in green, and sulfur atoms in yellow. **(B)** Inhibition dose response curve (IC_50_) for wild-type hP2X_4_ inhibited by BAY-1797 is measured to be 220 ± 30 nM from TEVC experiments in *Xenopus* oocytes. The Y-axis describes currents normalized to the largest current evoked by the maximum concentration of ATP applied to each oocyte (50 μM). The reported IC_50_ and error bars represent the mean and standard deviation across triplicate experiments, respectively. **(C)** Magnified view of the red box from panel A showing the BAY-1797 binding pocket. Important hydrogen-bonding interactions between BAY-1797 and proximal protein residues are shown with the yellow dotted lines. **(D)** The top-down view of hP2X_4_ bound to BAY-1797 highlighting the three-fold symmetry binding mode of BAY-1797. Each molecule of BAY-1797 binds in close proximity (3.3 Å apart), near the axis of symmetry of the receptor. **(E)** Zoomed in view of the allosteric binding pocket rotated 45° from panel C. In addition to showing several important hydrogen bonds (yellow dotted lines), this view highlights how W84 forms an edge-to-face interaction with BAY-1797. Except for residue A297, the dark yellow colored protomer is hidden in panels C and E to improve visual clarity.

BAY-1797 is bound in the canonical allosteric binding site of P2X_4_, located in the extracellular domain near the top of the receptor, and formed by upper body domains from neighboring protomers (Fig. 5, S3, S9) (*24*). At a resolution of 2.6 Å, the ligand density and density of the residues in the surrounding pocket are well-resolved, allowing us to model the position and orientation of BAY-1797 unambiguously (Fig. S8B). Ligand binding is mediated via hydrogen bonding interactions from several adjacent residues. The nitrogen of the sulfonamide moiety forms hydrogen bonds with a water molecule and the backbone carbonyl oxygen of A87 and V90 (distances of 3.4 Å, 3.1 Å, and 2.7 Å, respectively) (Fig. 5C,E, Fig. S8C). Additionally, the carbonyl oxygen of A297 from the adjacent monomer forms a hydrogen bond with the sulfonamide nitrogen (3.4 Å) (Fig. 5C,E). The amide in the phenylacetamide moiety of BAY-1797 forms hydrogen bonds with the sidechain amine of K298 and the backbone carbonyl oxygen of D88 (both at distances of 2.8 Å) (Fig. 5C,E). Aromatic interactions are also present: W84 forms an edge-to-face interaction with the benzyl group, with a centroid distance of 4.9 Å (Fig. 5C,E). Finally, the three molecules of BAY-1797 are close together, reaching within 3.3 Å of each neighboring, symmetry-related molecule (Fig. 5D).

In comparison with the published structure of zfP2X_4_ bound to BAY-1797, the allosteric ligand-binding pocket in hP2X_4_ is mostly conserved (Fig. 6A). The position of the allosteric pocket and the residues lining the pocket are homologous between hP2X_4_ and zfP2X_4_, and the overall orientation of the ligand, when bound in the pocket, is generally similar: the chlorophenoxy moiety is pointing inward toward the center of the receptor and the phenylacetamide group is pointing outward from the receptor (Fig. 6). However, there are several significant differences in the ligand binding pose and key receptor-ligand interactions that are ortholog-specific. In hP2X_4_, the sulfonamide moiety is rotated by 10° and inserted 0.9 Å deeper into the receptor (Fig. 6B,C). In zfP2X_4_, the chlorophenoxy group of BAY-1797 interacts edge-to-face with residue F299 at a distance of 6.4 Å. In hP2X_4_, however, this ligand pose would cause the chlorine atom of the chlorophenoxy group to clash with F296 in hP2X_4_; as a result, the chlorophenoxy moiety undergoes an 81° rotation about the horizontal x-axis to prevent contact with F296 but instead forms a “base stacking” interaction at a distance of 5 Å (Fig. 6C). In addition, the phenylacetamide group of BAY-1797 is rotated 42° off the receptor axis of symmetry, likely caused by the replacement of M108 in zfP2X_4_ with V105 in hP2X_4_, where the bulkier sidechain in the zebrafish ortholog causes the positional shift (Fig. 6D). Finally, because of the difference in sidechain orientation of K298 (K301 in zfP2X_4_), BAY-1797 forms hydrogen bonds with K298 in hP2X_4_, replacing the cation-π interactions with K301 that are present in zfP2X_4_ (Fig. 6B). A top-down view of the allosteric pocket highlights the different BAY-1797 poses between receptor orthologs (Fig. 6E). These significant differences in pose likely reflect inherent structural differences between the human and zebrafish orthologs of P2X_4_.

**Fig. 6.**
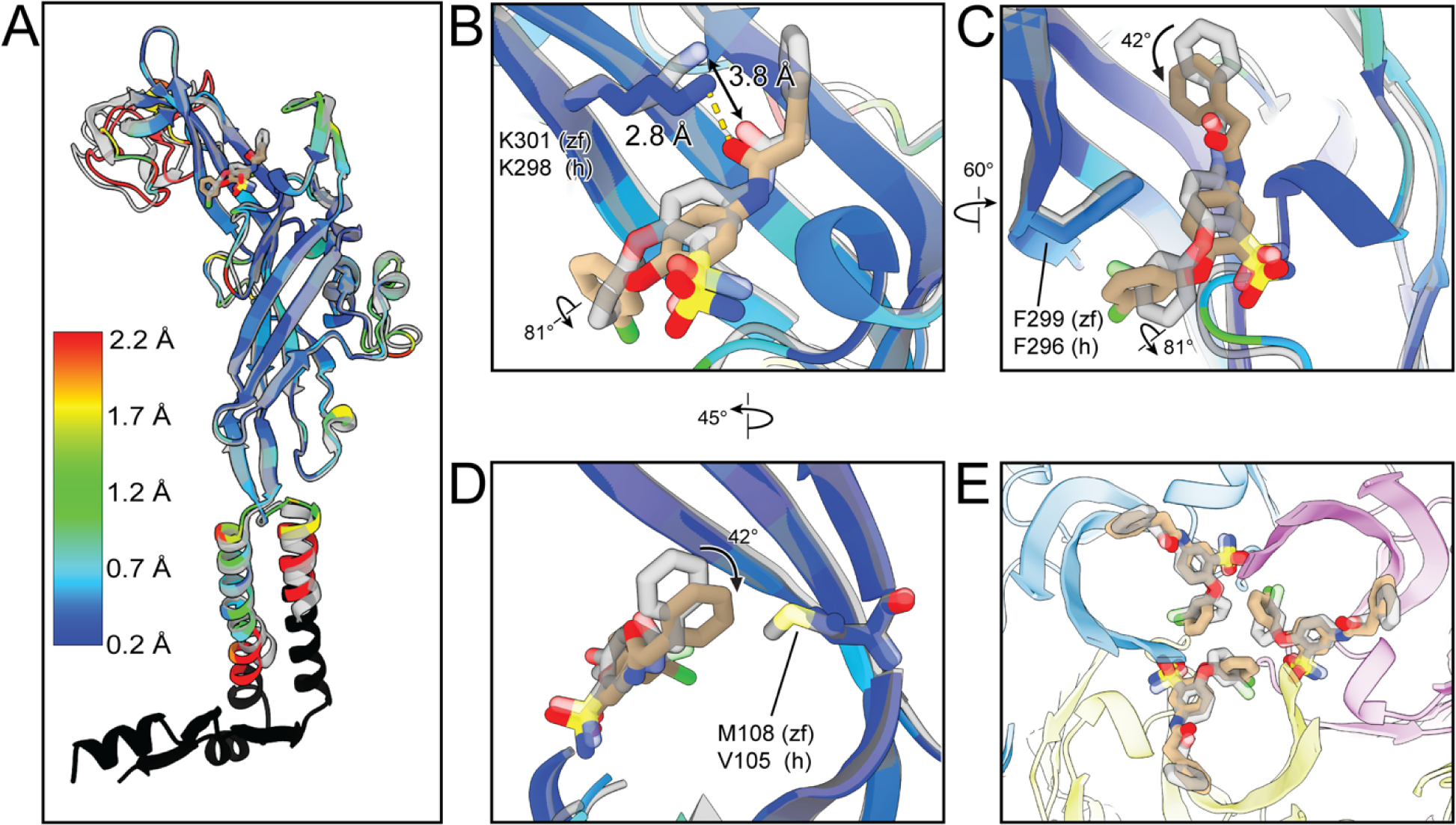
BAY-1797 binds to the allosteric binding site in hP2X_4_ with a different pose than zfP2X_4_. **(A)** Structural alignment of hP2X_4_ (ligand in tan, peptide colored by RMSD, with null values in black) and zfP2X_4_ (transparent grey) in complex with BAY-1797. The RMSD is color-coded to highlight ortholog-specific differences in structure. **(B)** Magnified view of the allosteric binding pocket in the same view as panel A highlighting the difference in pose for BAY-1797 when bound to hP2X_4_ compared to zfP2X_4_. In hP2X_4_, K298 is closer to BAY-1797, forming a hydrogen bond with the amide carbonyl (2.8 Å). In zfP2X_4_, the homologous residue K298 interacts with the ligand differently – a longer distance (3.8 Å) allows for a cation-π interaction only. **(C)** Magnified view of the allosteric pocket, rotated 60° from panel B, showing how a positional difference of F296 (F299 in zebrafish) affects the orientation of the cholorophenoxy group. **(D)** Magnified view of the allosteric pocket, rotated 45° from panel B, comparing the sidechains of V105 (human) and M108 (zebrafish) at the same structural position, where the smaller valine sidechain allows the phenylacetamide group to rotate 42° closer to the sidechain in hP2X_4_. **(E)** Top-down view of the allosteric ligand binding site showing how three molecules of BAY-1797 are close together in the binding pocket, reaching within 3.3 Å of each neighboring, symmetry-related molecule. In panel E, each protomer of hP2X_4_ is colored differently (light blue, light purple, and light yellow). For every panel in this figure, both molecules of BAY-1797 are colored by atom (see Fig. 5). The carbon atoms of the BAY-1797 molecule in the hP2X_4_ structure are colored tan. The carbon atoms of the BAY-1797 molecule in the zfP2X_4_ structure are light grey with all atoms transparent.

## Discussion

The structural and functional data generated from studying the full-length wild-type hP2X_4_ receptor provide key insights into subtype-specific structural features that play significant roles in P2XR desensitization (Fig. 7). (1) The cytoplasmic cap is already formed in the apo closed state of hP2X_4,_ prior to the formation of an open state, and then disassembles on the transition to the ATP-bound desensitized state. This finding redefines the helical recoil model of P2XR desensitization for slow-desensitizing subtypes, distinguishing it from fast and non-desensitizing P2XRs. (2) We identify putative lipid-binding sites in the cytoplasmic cap of hP2X_4_ that modulate receptor gating by affecting the rate of desensitization. (3) The hP2X_4_ receptor is significantly post-translationally modified, when expressed in mammalian cells, with extracellular glycosylation and cytoplasmic palmitoylation. (4) We highlight how binding of the noncompetitive antagonist, BAY-1797, does not impact the TM architecture, the presence of the cytoplasmic cap, or the presence of putative lipid-binding sites in the cytoplasmic domain relative to the apo closed state, but occupies the allosteric pocket of hP2X_4_ in an ortholog-specific pose.

**Fig. 7.**
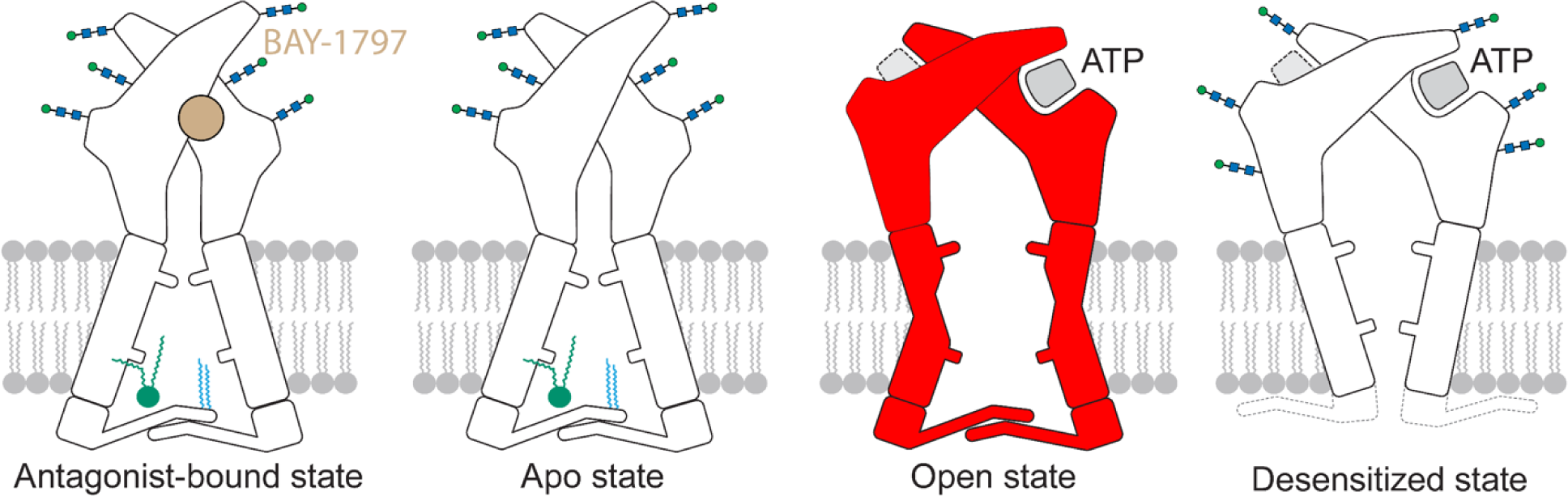
The gating cycle of the hP2X_4_ receptor redefines the helical recoil model of receptor desensitization for slow-desensitizing P2X receptors. Cartoon summarizing the gating mechanism for the hP2X_4_ receptor highlighting the presence of the cytoplasmic cap in the apo closed state, containing a putative lipid-binding site and palmitoylated residues. The three new conformational states from this manuscript are shown in white and the one conformational state extrapolated from a structure of the hP2X_3_ receptor is shown in red. In the redefined ‘helical recoil’ model for the slow-desensitizing P2X_4_ receptor, the cytoplasmic cap is present in the apo closed state – it does not form during the transition from the apo closed state to the ATP-bound open state, as was previously shown for the fast-desensitizing hP2X_3_ receptor. The stability of the cytoplasmic cap, modulated by putative lipid-binding sites, sets the lifetime of the ATP-bound open state. The disassembly of the cytoplasmic cap initiates the conformational change to the ATP-bound desensitized state. The overall architecture of the BAY-1797-bound inhibited state structure is very similar to the apo closed state structure – the binding of BAY-1797 does not impact the TM domains, the location of the gate, the presence of the cytoplasmic cap, or the presence of putative lipid-binding sites in the cytoplasmic domain.

Our full-length wild-type structures of the hP2X_4_ receptor create a paradigm shift in the helical recoil model of P2XR desensitization to account for slow-desensitizing subtypes (Fig. 7). Similar to the P2X_7_ receptor, the apo closed state of hP2X_4_ has an assembled cytoplasmic cap (Fig 1A, 2A). However, unlike the P2X_7_ receptor, the cytoplasmic cap of hP2X_4_ is not permanently stabilized by palmitoyl groups, as P2X_4_ undergoes slow desensitization (Fig. 2C, Fig. 3F) (*18*). On the other hand, the P2X_4_ receptor follows a similar gating mechanism to the P2X_3_ receptor, whereby the cytoplasmic cap is disassembled in the ATP-bound desensitized state (Fig. 4A). Thus, the P2X_4_ receptor can be thought of as an intermediate receptor, where its gating mechanisms are a blend of fast and non-desensitizing P2XR subtypes.

The structural basis for slow desensitization in the hP2X_4_ receptor originates from the stability of the cytoplasmic cap, modulated by inter-protomer residue interactions and putative lipid-binding sites. Importantly, the cytoplasmic cap was not resolved in any previous structural study of zfP2X_4_, likely due to two major distinctions in experimental methods (*22*–*24*). First, the terminal cytoplasmic residues that are key to maintaining cap-forming interactions were removed within some of the truncated zebrafish receptor constructs. We and others have shown that mutation or truncations of residues within the N- or C-termini impact the function of P2XRs (*22*, *34*–*40*). Second, the detergent conditions (for crystallization) or amphipols (for cryo-EM) may not have provided a suitable environment to stabilize the cytoplasmic cap. To this point, our structures identify two detergent-binding sites in the cytoplasmic domain of hP2X_4_ that are key to its stabilization (Fig. 3A).

Previous studies have demonstrated the importance of phosphoinositides for P2X_4_ receptor activation through direct interactions in the C-terminus (*7*, *32*). Thus, there is precedence to suggest that the two DDM molecules present in our models of hP2X_4_ might reflect endogenous lipid-binding sites in the native environment. Indeed, we show that mutations in the cytoplasmic domain of hP2X_4_ aimed at reducing the ability to bind putative lipids, based on interactions with the DDM detergent in our structure, resulted in faster rates of desensitization *in vivo,* likely through reducing the stability of the cytoplasmic cap (Fig. 3F). The first detergent molecule binds vertically in the same direction as the TMs and directly interacts with residues R368 and R33, two positively charged residues that could interact strongly with a negatively charged head group of a native phospholipid (Fig. 1A, 3A,B). The second detergent binds perpendicular to the TMs and interacts with residue D16 and D354 (Fig. 3C). Previous tryptophan scanning studies have identified that mutation of D354 abrogates receptor function while not affecting trafficking (*41*). Similarly, we find that mutation of R33 significantly diminishes ATP-induced activity. The interactions of R33 and D354 with a native lipid may be necessary contacts for native function of the receptor; while the residues R368 and D16 might provide interactions important for the characteristic slow-desensitization kinetics of hP2X_4_. We speculate that one native lipid could occupy both detergent-binding sites, with one tail positioned vertically and parallel to the TMs and the other tail positioned horizontally, perpendicular to the TMs. Additional studies are needed to identify the native lipid and determine how these putative protein-lipid interactions within the cytoplasmic cap of P2X_4_ affect the rate of receptor desensitization.

The hP2X_4_ receptor has a significant amount of post-translational modifications. It is highly glycosylated when expressed in mammalian cells, more so than any other structurally characterized P2XR (*16*, *18*, *22*–*25*). The uniquely high number of glycosylated residues present on P2X_4_ was demonstrated to be necessary for membrane localization and receptor hydrophilicity (*42*, *43*). Furthermore, the current model of P2X_4_ recycling postulates that P2X_4_ is internalized in acidic lysosomal compartments for re-sensitization, where glycosylation prevents receptor degradation (*43*). In addition to glycosylation, we discovered that the cytoplasmic domain of hP2X_4_ is palmitoylated. However, these post-translational modifications do not affect receptor desensitization and their roles are yet to be determined (Fig. 2C, S6). Further, it is unclear if the heterogenous palmitoylation we observe indicates redundancy between palmitoylation sites or if protein overexpression for MS studies has prevented uniform palmitoylation. Additional work should be conducted to elucidate the functional consequences, if any, of hP2X_4_ palmitoylation.

The structure of hP2X_4_ in an inhibitor-bound state is highly similar to the apo state structure of hP2X_4_, which agrees with previous structural studies on inhibited P2X receptors (Fig. 1A, 4A, 6) (*24*, *25*, *44*). Ligand binding in the allosteric site has been postulated to prevent receptor transition to the open state, rather than affect ATP binding (*25*). In agreement with this model, the cytoplasmic cap remains ordered and, except for an expanded allosteric pocket to accommodate inhibitor binding, the overall receptor architecture is not significantly altered (Fig. 1A, Fig. 4A). While the potency of BAY-1797 inhibition is quite similar between human and zebrafish subtypes of P2X_4_, there are significant differences in the local allosteric site and the pose of the ligand in the pocket. The specific pose of the BAY-1797 bound to the human subtype provides more accurate information for future drug modification strategies. For example, the position of the phenylacetamide ring in the human ortholog allows space for a meta substitution on the phenyl group which could provide more specificity. Further, the orientation of the chlorophenoxy moieties in the human ortholog could allow engineering of functional groups that form specific inter-ligand interactions to increase affinity. Despite the structural and sequence similarity between the two receptor orthologs, the observed differences in ligand pose are evidence that future studies should focus on the human subtype for most relevance with respect to structure-based drug development efforts. Altogether, our data provides the structural basis of allosteric antagonism for the human ortholog and elucidates the key structural determinants of gating kinetics for slow-desensitizing P2XR subtypes, thereby redefining the helical recoil model.

## Materials and Methods

### Cell Lines

SF9 cells were cultured in SF-900 III SFM (Fisher Scientific) at 27°C and used for expression of Baculovirus (cells of female origin). Receptor was expressed in HEK293 GNTI-cells that were cultured in Gibco Freestyle 293 Expression Medium (Fisher Scientific) at 37°C, supplemented with 2% v/v fetal bovine serum (FBS) (HEK293 cells of female origin).

Unfertilized *Xenopus* laevis oocytes were purchased through Ecocyte Biosciences and kept at 18°C until injection.

### Receptor Constructs

The full-length, wild-type hP2X_4_ receptor construct used for structure determination contains a C-terminal GFP, a 3C protease site, and a histidine affinity tag for purification. No mutations or truncations were made to the receptor for structure determination. For electrophysiology experiments, the hP2X_4_-WT construct is not genetically mutated and does not contain GFP, protease sites, or affinity tags. Mutations were performed using QuikChange XL mutagenesis kits (Agilent Technologies) from this wild-type construct. The following set of mutants were generated for electrophysiology experiments: C4A/C5A, C4A/C5A/C360A, D16A, R33A, R368A, and R368A/Y359F.

### Receptor Expression and Purification

Full-length wild-type hP2X_4_ was expressed using identical protocols previously outlined (16, 18). Briefly, HEK293 GNTI-cells were grown to a sufficient density before being infected with high-titer P2 hP2X_4_ BacMam virus. Sodium butyrate (10 mM) was added after overnight incubation at 37 °C, before moving the cells to 30 °C to express for 2 additional days. After sufficient expression, cells were harvested with PBS (137 mM NaCl, 2.7 mM KCl, 8 mM Na_2_HPO_4_, and 2 mM KH_2_PO_4_), resuspended in TBS (50 mM Tris pH 8.0, 150 mM NaCl) with protease inhibitors (1 mM PMSF, 0.05 mg/mL aprotinin, 2 mg/mL Pepstatin A, 2 mg/mL leupeptin), and lysed by sonication. Cellular debris was removed by a low-speed centrifugation, after which membranes were isolated following a high-speed centrifugation, frozen, and stored at –80 °C until use.

Membranes were thawed, resuspended in TBS buffer + 15% glycerol, and dounced until homogenous. Next, the homogenized membranes were solubilized in 40 mM n-Dodecyl β-D-maltoside (DDM) and 8 mM cholesteryl hemisuccinate (CHS) for 2 hours and ultracentrifugation performed to isolate the soluble fraction, which was then bound to TALON cobalt resin for 1 hour in the presence of 10 mM imidazole. The resin was packed into an XK-16 column and the protein complex was eluted in buffer (TBS, 5% glycerol, 1 mM DDM, 0.2 mM CHS) containing a gradient of imidazole (10 to 250 mM). Protein-containing fractions were collected, concentrated, and cleaved by HRV 3-C protease overnight at 4 °C to remove the GFP-His tag. Digested protein was cleared by ultracentrifugation and then run on a Superdex 200 10/300 GL column in SEC buffer (20 mM HEPES pH 7.0, 100 mM NaCl, 0.5 mM DDM) to separate trimeric receptor from cleaved GFP. After performing fluorescence size exclusion chromatography, size exclusion chromatography (SEC) fractions containing high-quality protein receptor were pooled and concentrated for cryo-EM grid preparation.

### Electron Microscopy Sample Preparation

Purified hP2X_4_ (2.5 μL, 5.5 mg/mL) was applied to a glow-discharged (15 mA, 1 min) Quantifoil R1.2/1.3 300 mesh gold holey carbon grid and blotted for 1.5 seconds under 100% humidity at 6 °C. The grids were then flash frozen in liquid ethane using a FEI Vitrobot Mark IV before being clipped and sent for screening and data collection.

### Electron Microscopy Data Collection and Processing

The two datasets for hP2X_4_ were collected on Titan Krios Microscopes (FEI) operated at 300 kV and at a nominal magnification of 130,000x. All data was acquired at the Pacific Center for Cryo-EM (PNCC) using Gatan K3 direct-electron detectors and energy filters (Gatan Image Filter). Movies for the combined apo closed and ATP-bound desensitized dataset were collected in super-resolution mode with a physical pixel size corresponding to 0.648 Å/pixel, a defocus range of -0.9 to -1.5 µm, and a total dose of 43 e^-^/ Å^2^. Movies for the BAY-1797-bound inhibited dataset were collected hardware-binned at a physical pixel size of 0.647 Å/pixel, a defocus range of -0.7 to -1.4 µm, and a total dose of 45 e^-^/ Å^2^. Each dataset utilized ‘multi-shot’ and ‘multi-hole’ collection schemes driven by serialEM to maximize high-throughput data collection (46).

### Electron Microscopy Data Processing

Movies were patch motion corrected in cryoSPARC and micrographs generated at the physical pixel size (Table S1) (47). Contrast transfer function (CTF) parameters were estimated in cryoSPARC and micrographs were manually curated (47). Particles were picked using 2D-templates generated from a 3D reconstruction, inspected to remove bad picks, and extracted at a 2x binned pixel size. Particles were then directly sent to 3D-classification, skipping 2D classification, using a combination of ab initio jobs to generate 3D volumes followed by heterogenous refinements. After a final particle stack was reached, particles were re-extracted at the physical pixel size and sent to a final non-uniform refinement, using local and global CTF refinement, which generated the consensus reconstruction. Some maps were sharpened in Scipion 3 using localdeblur sharpening from Xmipp3 (48, 49).

### Model Building and Structure Determination

Homology models for the apo closed and ATP-bound desensitized state of hP2X_4_ were generated from SWISS-MODEL using the apo closed state of zfP2X_4_ and the ATP-bound desensitized state of hP2X_3_ (PDB codes: 4DW0 and 5SVL, respectively) (50). The cytoplasmic cap of hP2X_4_ in the apo closed state was built de novo. The initial structures were then built in COOT and iteratively refined in PHENIX, using CIF files generated with eLBOW (51–53). Limited glycosylations and detergent molecules were modeled when justified by density. In some models, residues side chains are not modeled beyond the α-carbon if density was not supportive. The BAY-1797-bound inhibited state model was generated from the finalized apo closed state model of hP2X_4_. All models were evaluated by MolProbity to check quality (Table S1) (54).

### Two-Electrode Voltage Clamp

#### Preparation of Oocytes expressing hP2X_4_

Defolliculated *Xenopus* oocytes were ordered from Ecocyte and resuspended in Barth’s solution (88 mM NaCl, 1 mM KCl, 0.82 mM MgSO_4_, 0.33 mM Ca(NO_3_)_2_*4H_2_O, 0.41 mM CaCl_2_*2H_2_O, 2.4 mM NaHCO_3_, 5 mM HEPES) supplemented with amikacin (250 mg/L) and gentamycin (150 mg/L). Oocytes were then injected with 50 nL of 200 ng/µL full-length wild-type or mutant hP2X_4_ mRNA synthesized from pCDNA3.1x using the mMessage mMachine T7 Ultra Kit (Invitrogen). The oocytes were incubated at 18 °C for 2 days to allow sufficient receptor expression. Sutter filamented glass (length: 10 cm, inner diameter: 0.69 mm, outer diameter: 1.2 mm) was filled with 3 M KCl and used to impale the injected oocytes and clamp at a holding voltage of –60 mV.

#### TEVC Recordings

Data acquisition was performed using the Ooctye Clamp OC-725C amplifier and pClamp 8.2 software. Gravity fed RSC-200 Rapid Solution Changer was used to apply buffer at a rate of ∼5 mL/min. Sutter filamented glass were prepared at 10 cm in length with an inner diameter of 0.69 mm and an outer diameter of 1.2 mm to impale oocytes. The holding voltage was clamped at -60 mV. Experiments were recorded in buffer containing 10 mM HEPES (pH 7.4), 140 mM NaCl, 5 mM KCl, 2 mM CaCl_2_, 2 mM MgCl_2_, and 10 mM glucose.

#### Dose Response (EC_50_) Experiments

Excitatory responses to a dilution series of ATP consisting of 520 µM, 300 µM, 100 µM, 33.3 µM, 11.1 µM, 3.7 µM, 1.2 µM, and 0.4 µM were tested. Oocytes expressing hP2X_4_ were briefly exposed to 300 µM ATP several times every 60 seconds until the excitatory responses reached a constant maximum amplitude, at which point the ATP test concentration was applied. Each evoked ATP response was normalized to the preceding 300 µM ATP signal. Each ATP concentration was repeated in triplicate across different oocytes. An EC_50_ dose-response curve was generated in Prism 9 by fitting the data to a nonlinear regression named: “EC_50_, x is concentration.” All currents were normalized to a preceding current evoked by an application of 300 µM ATP. The three discrete trials for each condition were used to generate an average which was reported plus or minus its standard deviation.

#### Inhibition Dose Response (IC_50_) Experiments

Inhibitory responses to a dilution series of BAY-1797 consisting of 20 µM, 6.66 µM, 2.22 µM, 740 nM, 247 nM, 82 nM, 27 nM, and 9 nM were examined. As with the agonist dose response experiment, 50 µM ATP was applied every two minutes to obtain a constant evoked current. The oocyte was then washed with buffer and antagonist for two minutes, followed by co-application of the same test concentration with 50 µM ATP. The antagonized signal was then normalized against the initial 50 µM ATP excitatory signal. Each test concentration was performed in triplicate across different oocytes. The data was then fit to a nonlinear regression named "[inhibitor] vs. response – Variable slope (four parameters)" in Prism 9 to produce a sigmoidal curve and determine an IC_50_ value. The three discrete trials for each condition were used to generate an average which was reported plus or minus its standard deviation.

#### Mutant Receptor Kinetics Experiments

Oocytes expressing mutant hP2X_4_ (C4A/C5A, C4A/C5A/C360A, D16, R33A, R368, Y359F/R368A) were repeatedly exposed to 50 µM ATP for one minute followed by a two-minute buffer wash. The resulting traces were then normalized to WT current traces to observe differences in receptor kinetics. Desensitization lifetimes (*τ* values) were calculated by performing a time-weighted average of *τ*_fast_ and *τ*_slow_ values after fitting the desensitization traces with a two-phase decay function. The magnitude of the currents for the palmitoylation mutants (C4A/C5A, C4A/C5A/C360A) was very similar to that of wild-type hP2X_4_. However, the other mutants (D16A, R368A, and Y359F/R368A) had reduced expression (current amplitudes between 400-800 nA, 300-750 nA, and 200-400 nA, respectively) compared to wild-type hP2X_4_ in oocytes (current amplitudes between 1 to 3 µA). The R33A mutation was impacted more severely as many oocytes featured essentially no expression or low signal (below 100 nA).

### Mass spectrometry for detection of palmitoylation sites

Samples were dried by centrifugal evaporation and reconstituted with 5% SDS, 8 M Urea, 100 mM Glycine (pH 7.5). Cysteines were chemically modified by the addition of tris(2-carboxyethyl)phosphine (TCEP) followed by incubation with methyl methanethiosulfonate (MMTS). Samples were acidified by the addition of phosphoric acid and SDS protein extraction buffer (90% methanol, 100 mM TEAB, pH 8) was added. Samples were transferred to S-trap micro columns (Protifi, Farmingdale, NY) and washed with 90% methanol, 100 mM TEAB. S-trap columns containing the bound sample proteins were digested with sequencing grade modified trypsin (Promega, Fitchburg, WI, Cat # V5111). Upon completion of digestion, peptides were eluted and then dried by vacuum centrifugation. Peptide concentrations were determined using a Pierce Quantitative Colorimetric Peptide Assay (ThermoFisher Scientific, Cat # 23275) before LC-MS/MS analysis. Two µg injections of peptides were then analyzed using a Q-Exactive HF mass spectrometer (ThermoScientific) coupled to an UltiMate 3000 RSLCnano liquid chromatograph instrument in the OHSU Proteomics Shared Resource Core Facility. Electrospray ionization was performed using a EasySpray nano source and 1.0 kV source voltage. Xcalibur version 4.0 was used to control the system. Samples were applied at 5 µl/min for 5 min to an Acclaim PepMap 100 μm x 2 cm NanoViper C18 peptide trap using a loading pump and 2% acetonitrile (ACN), 0.1% formic acid solvent. Peptides were then separated using a 7.5–30% ACN gradient over 60 min, and 30-98% ACN gradient over 2 min in a mobile phase containing 0.1% formic acid at a 300 nl/min flow rate. Survey mass spectra were acquired over m/z 375−1400 at 120,000 resolution (m/z 200), maximum ion time of 50 ms, and AGC target of 3x106. Data-dependent acquisition selected the top 10 most abundant precursor ions for MS2 using HCD fragmentation, an isolation width of 1.2 m/z, normalized collision energy of 30, resolution of 30,000, maximum ion time of 100 ms, minimum AGC of 5x103, dynamic exclusion set to auto, and charge state selection from of +2 to +7. Raw MS data files searched against the UniProt human proteome database from Dec 2023 using SequestHT in the Protein Discoverer 2.5 suite. Differential modifications included removal of protein N-terminal methionine and acetylation of the resulting N-termini, as well as palmitoylation and methylthio dynamic modifications of cysteines. A precursor mass tolerance of 20 ppm and fragment ion tolerance of 0.2 Daltons was used. Results were filtered using Percolator using a reversed sequence strategy to measure peptide false discovery rates and only peptides with q scores < 0.01 were excepted. The analysis assigned 82.99% of the human P2X_4_ receptor sequence.

## Acknowledgments

We thank O. Davulcu, C. Yoshioka, and C. López at PNCC for access and microscopy assistance. We thank L. Anson for comments on the manuscript. We thank L. David, A. Reddy, and K. Zientek at the proteomics core at Oregon Health & Science University (OHSU). Electron microscopy grid screening was performed at the Multiscale Microscopy Core and mass spectrometry experiments were performed at the Proteomics core within OHSU.

## Funding

A portion of this research was supported by NIH grant U24GM129547 and performed at the PNCC at OHSU and accessed through EMSL (grid.436923.9), a DOE Office of Science User Facility sponsored by the Office of Biological and Environmental Research.

National Institute of Health grant R00HL138129 (SEM)

National Institute of Health grant DP2GM149551 (SEM)

American Heart Association grant 24PRE1195450 (ACO)

## Author contributions

SEM designed the project. HS and IAD performed sample preparation for cryo-EM. ACO performed the cryo-EM data collection. HS and ACO analyzed the cryo-EM data. HS and ACO built the cryo-EM models. IAD performed and analyzed the electrophysiological experiments. All authors wrote and edited the manuscript.

## Competing interests

Authors declare that they have no competing interests.

## Data and materials availability

All data are available in the main text or the supplementary materials. Cryo-EM density maps for the full-length wild-type hP2X_4_ receptor in the apo closed, ATP-bound desensitized, and BAY-1797-bound inhibited states have been deposited in the EM Database under the EMDB accession codes: EMD-44799, EMD-45177, and EMD-44800, respectively. The maps within these depositions include both half maps, sharpened/unsharpened maps, refinement masks, and any locally sharpened maps that helped with model building. The corresponding coordinates for the structures have been deposited in Protein Data Bank under the PDB accession codes: 9BQH, 9C48, and 9BQI, respectively.

## Notes

### Competing Interest Statement

The authors have declared no competing interest.

