## Supplemental Files for "Human P2X4 receptor gating is modulated by a stable cytoplasmic cap and a unique allosteric pocket"

**This PDF file includes:**

Figs. S1 to S9
Table S1

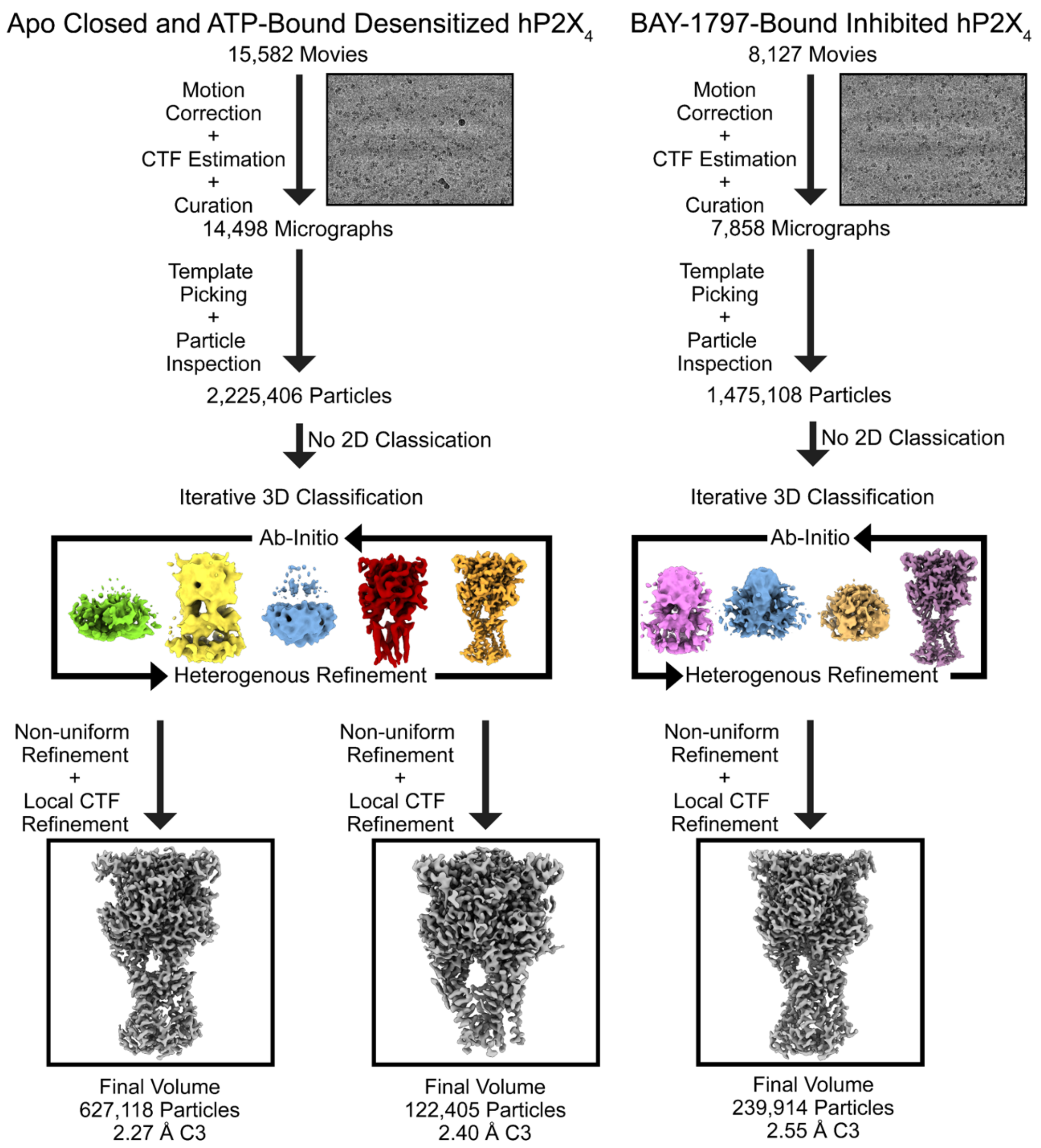


**Fig. S1. Cryo-EM processing pipeline for hP2X4 reconstructions.** Image processing pipeline for hP2X4 structures in the apo closed, ATP-bound desensitized, and BAY-1797-bound inhibited states using CryoSPARC (47). Movies were motion corrected, CTF parameters estimated, and curated. After template picking, particles were inspected and extracted. Skipping 2D-classification, particles were passed directly to iterative 3D classification, alternating between ab-initio and heterogenous refinements, to obtain a final particle stack. Final particles were then subjected to further CTF and non-uniform refinements at the physical pixel size to generate the final reconstructions.

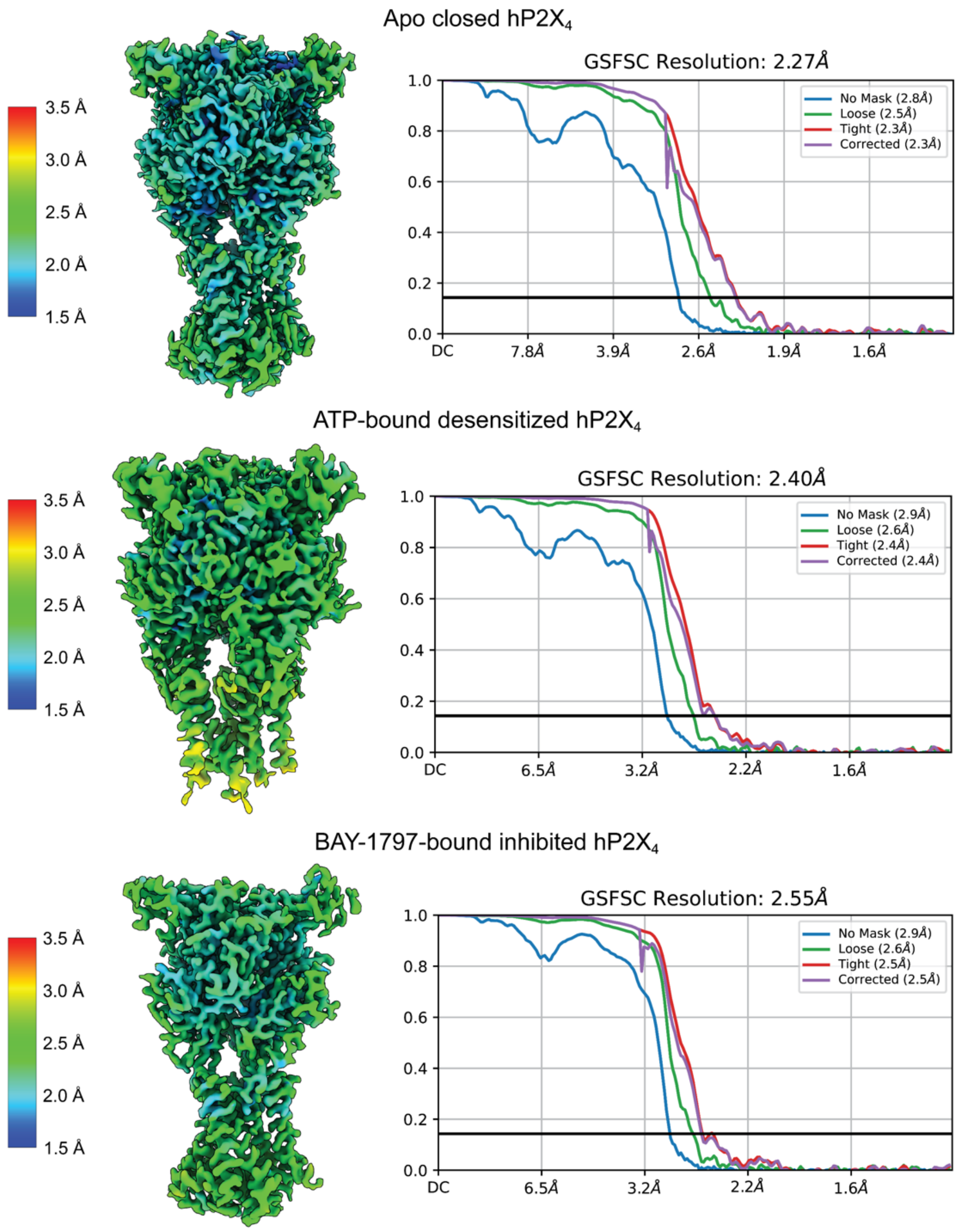


**Fig. S2. Local resolution maps and gold-standard FSC graphs for the structures of $hP2X_4$ in the apo closed state, the ATP-bound desensitized state, and the BAY-1797-bound inhibited state.** The resolutions stated were measured at an FSC threshold of 0.143. Local resolution estimations were provided at a range between 1.5 Å (blue) and 3.5 Å (red).

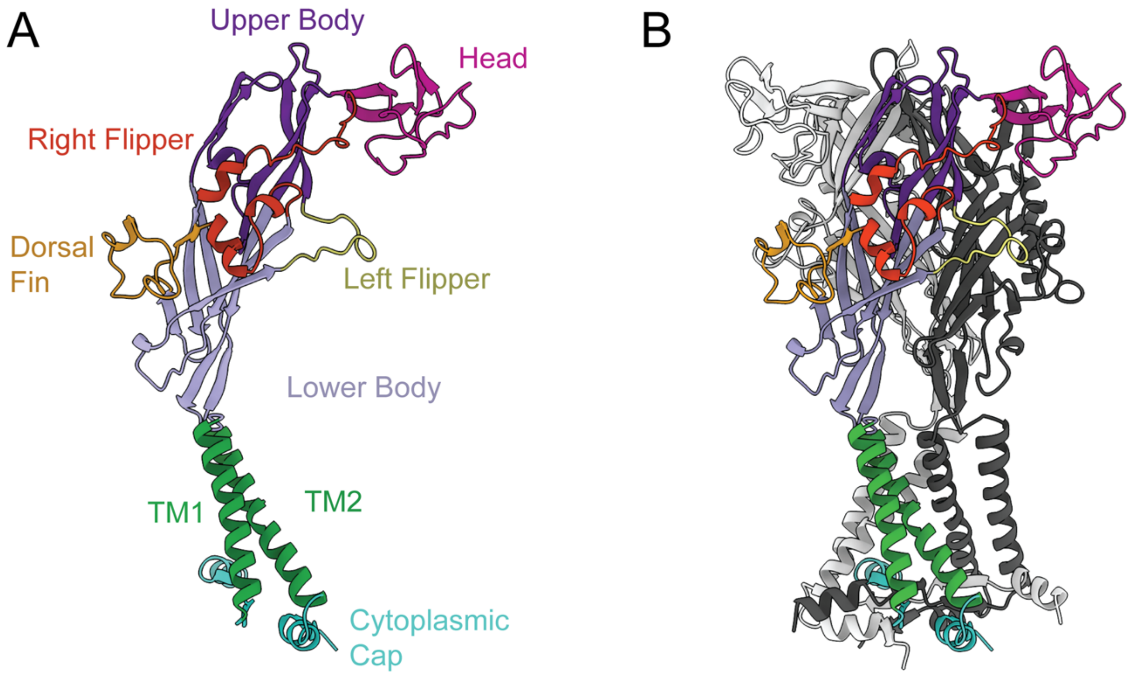


**Fig. S3. Naming of the purinergic receptor domains. (A and B)** Ribbon representation of one protomer **(A)** as well as the full trimeric receptor **(B)** for the apo closed state of $hP2X_4$ colored by domain architecture. For the full receptor, the second and third protomers are colored light grey and dark grey, respectively.

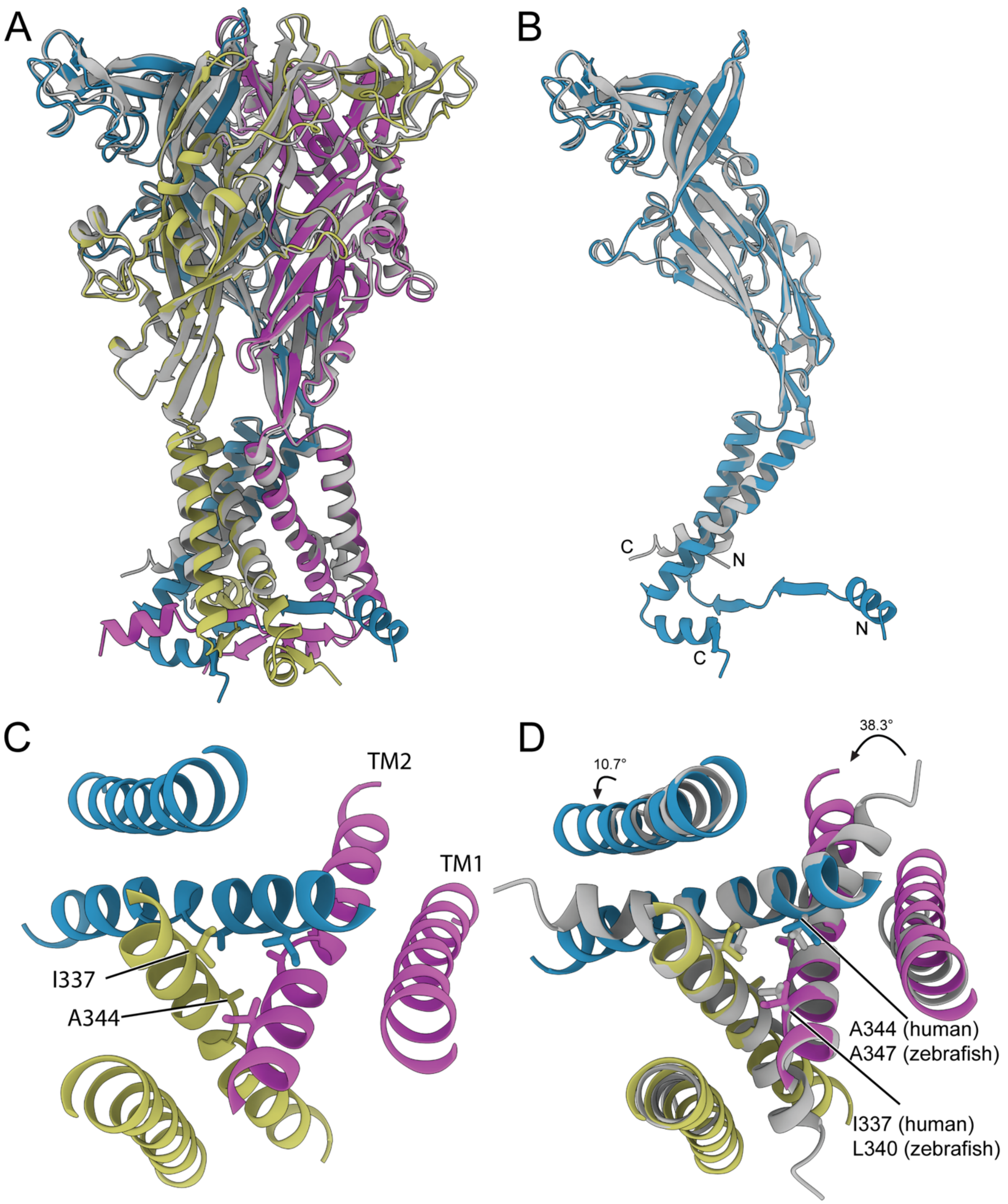


**Fig. S4. Structural alignment of $hP2X_4$ in the apo closed state to $zfP2X_4$ in the apo closed state. (A)** Overlayed apo closed state structure of $hP2X_4$ (protomers colored blue, purple and dark yellow) and the apo closed state structure of $zfP2X_4$ (light grey, PDB code 4DW0) highlighting the differences between the two orthologs. The largest differences come in the TM domain and the cytoplasmic domain where the full-length $hP2X_4$ structure is significantly more complete, containing TM domains that span the lipid bilayer with a cytoplasmic cap. **(B)** Ribbon representation of one subunit of hP2X4 in the apo closed state (blue) aligned with one subunit of zfP2X4 in the apo closed state (light grey). **(C)** Top-down view of the ion channel pore of the hP2X4 receptor in the apo closed state. The conductance pathway of hP2X4 is blocked at two locations: an initial constriction gate formed by I337 from each protomer (1.0 Å pore radius) and then a tighter constriction gate formed by A344 from each protomer (0.4 Å pore radius). **(D)** Top-down view of the overlayed apo closed state structure of hP2X4 (protomers colored blue, purple and dark yellow) and the apo closed state structure of zfP2X4 (light grey), highlighting the similarities in the pore architecture between the two orthologs. The two constriction gates of hP2X4 in the apo closed state (I337 and A344) align nearly perfectly with the gates of zfP2X4 in the apo closed state: L340 (0.7 Å pore radius) and A347 (0.7 Å pore radius). The TM domains of the two orthologs also differ where TM1 is rotated by 10.7° and TM2 is rotated by 38.3°.

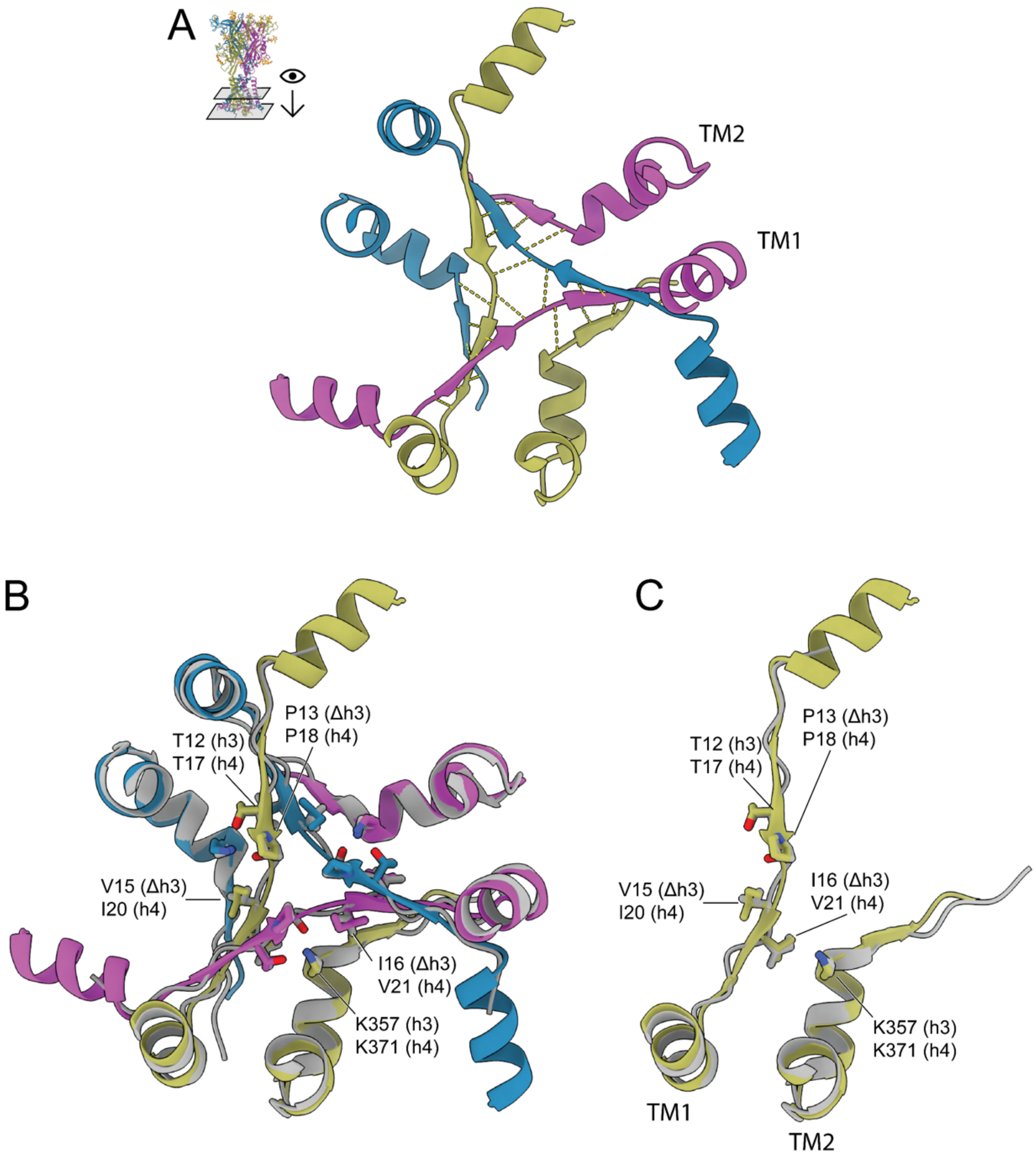


**Fig. S5. The cytoplasmic cap of hP2X$_4$ in the apo closed state highlighting extensive hydrogen bonding and homology with the ATP-bound open state of hP2X$_3$. (A)** Top-down view of the cytoplasmic cap of hP2X$_4$, with the hydrogen bonding interactions important for the cytoplasmic cap structure shown as dashed yellow lines. Each protomer of the receptor is a different color: blue, purple, and dark yellow. **(B)** Structural alignment of the cytoplasmic cap in the context of a full trimeric receptor of hP2X$_4$ in the apo closed state (protomers colored blue, purple and dark yellow) with the cytoplasmic cap in the context of a full trimeric receptor of chimeric hP2X$_3$ (light grey) in the ATP-bound open state. The ATP-bound open state structure of hP2X$_3$ was obtained using a P2X$_2$/P2X$_3$ chimeric construct containing three key mutations in the cytoplasmic cap: T13P, S15V, and V16I. The alignment was made between residues 13-33 and 356-372 of hP2X4 and residues 8-28 and 342-358 of chimeric $hP2X_3$ (PDB code: 5SVK). Residues found in wild-type $hP2X_4$, wild-type $hP2X_3$, and chimeric $hP2X_3$ are indicated as h4, h3, and Δh3, respectively. **(C)** Structural alignment of a single protomer of the cytoplasmic cap of $hP2X_4$ in the apo closed state (dark yellow) with a single protomer of the cytoplasmic cap of chimeric $hP2X_3$ in the ATP-bound open state (light grey).

A

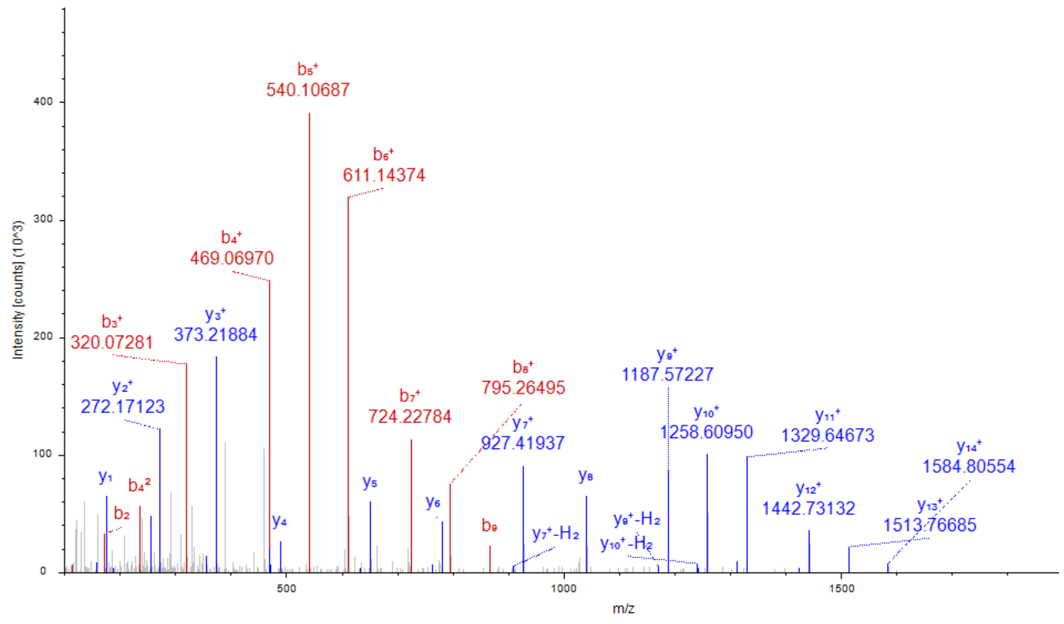

Intensity [counts] (10^3)
m/z
$b_5^+$ 540.10687
$b_6^+$ 611.14374
$b_4^+$ 469.06970
$b_3^+$ 320.07281
$y_3^+$ 373.21884
$y_2^+$ 272.17123
$b_7^+$ 724.22784
$b_8^+$ 795.26495
$y_7^+$ 927.41937
$y_9^+$ 1187.57227
$y_{10}^+$ 1258.60950
$y_{11}^+$ 1329.64673
$y_{12}^+$ 1442.73132
$y_{13}^+$ 1513.76685
$y_{14}^+$ 1584.80554
$y_1$
$b_4^2$
$b_2$
$y_4$
$y_5$
$y_6$
$b_9$
$y_7^+$-$H_2$
$y_8$
$y_9^+$-$H_2$
$y_{10}^+$-$H_2$


B

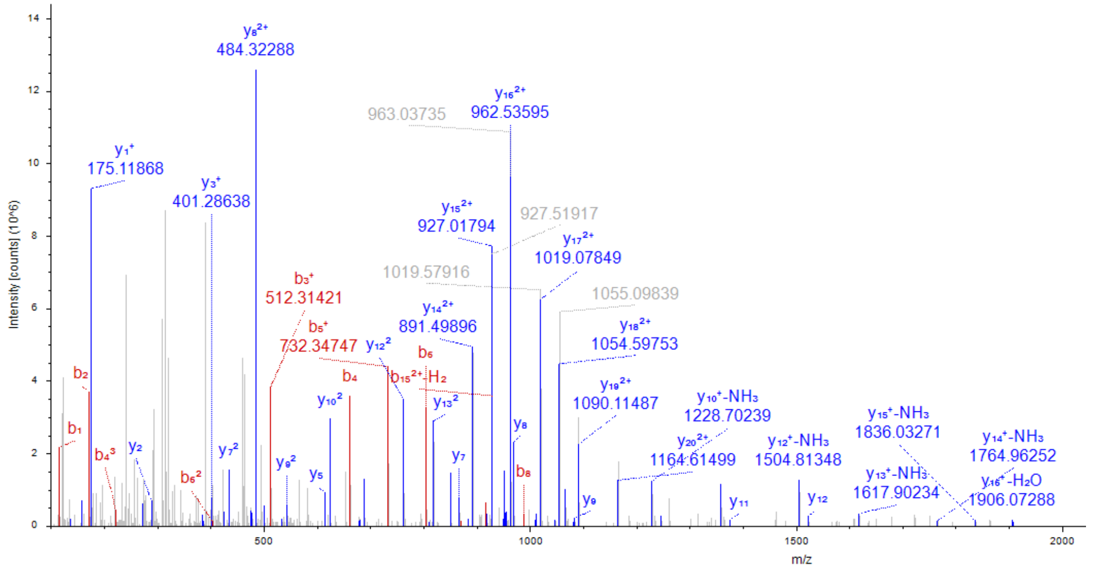

Intensity [counts] (10^6)
m/z
$y_8^{2+}$ 484.32288
963.03735
$y_{16}^{2+}$ 962.53595
$y_1^+$ 175.11868
$y_3^+$ 401.28638
$y_{15}^{2+}$ 927.01794
927.51917
$y_{17}^{2+}$ 1019.07849
1019.57916
1055.09839
$b_3^+$ 512.31421
$y_{14}^{2+}$ 891.49896
$b_5^+$ 732.34747
$y_{12}^2$
$y_{18}^{2+}$ 1054.59753
$b_6$
$b_4$
$b_{15}^{2+}$-$H_2$
$y_{19}^{2+}$ 1090.11487
$y_{10}^+$-$NH_3$ 1228.70239
$y_{15}^+$-$NH_3$ 1836.03271
$y_{14}^+$-$NH_3$ 1764.96252
$y_{10}^2$
$y_{13}^2$
$y_8$
$y_{20}^{2+}$ 1164.61499
$y_{12}^+$-$NH_3$ 1504.81348
$y_{13}^+$-$NH_3$ 1617.90234
$y_{16}^+$-$H_2O$ 1906.07288
$b_2$
$b_1$
$b_4^3$
$y_2$
$y_7^2$
$b_6^2$
$y_9^2$
$y_5$
$y_7$
$b_8$
$y_9$
$y_{11}$
$y_{12}$

C

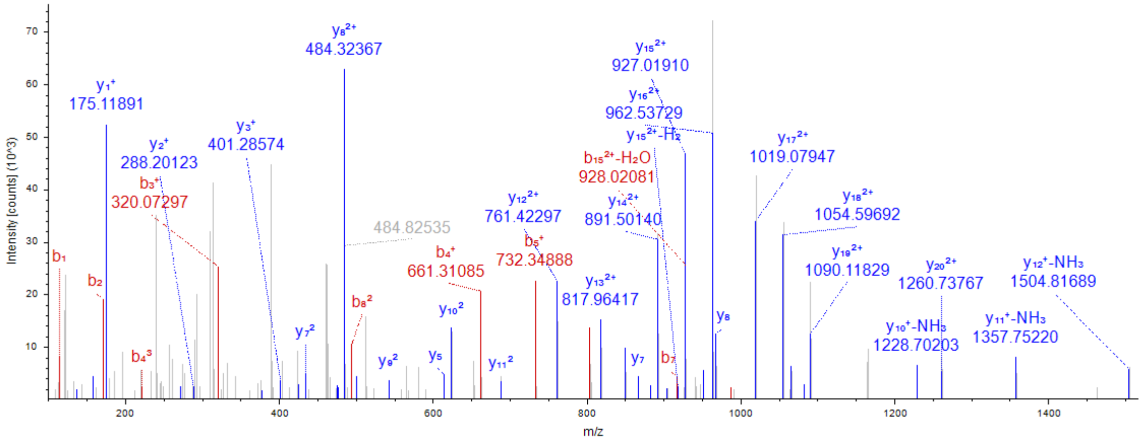


D

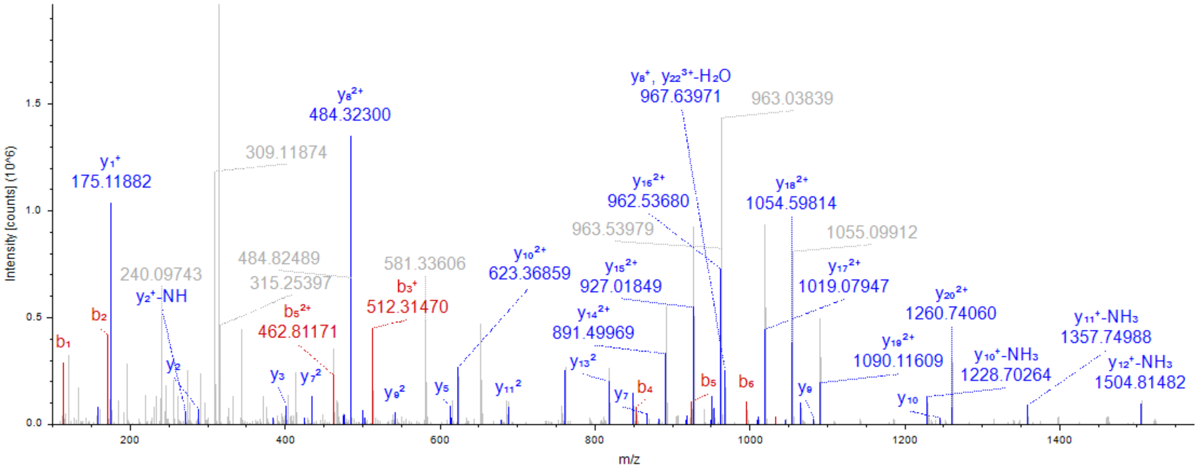


**Fig. S6: MS/MS spectra identification of modifications in the triply charged N-terminally acetylated peptide 1-23 of P2X4 (acetyl-AGCCAALAAFLFEYDTPRIVLIR). (A)** Non-palmitoylated version (precursor m/z= 883.43), identified by assigned b3+ (320.07) and b4+ (469.07) ions that contain methylthio (blocking agent) modifications at C3 and C4. **(B)** Single palmitoylated version (precursor m/z = 947.52), identified by assigned b3+ (512.31) and y202+ (1165.12) ions containing a palmitoylation at C3, and methylthio blocking group at C4. **(C)** Single palmitoylated version (precursor m/z = 947.52), identified by assigned b3+ (320.07) and y202+ (1261.73) ions containing a palmitoylation at C3, and methylthio blocking group at C4. Note: MS/MS spectra in B and C are identical, but assigned fragment ions somewhat differ due to location of the palmitoyl group at either C3 or C4, because both isoforms coelute from the LC column and fragment simultaneously. **(D)** Doubly palmitoylated version (precursor m/z = 1011.93) identified by assigned b3+ (512.31) and y202+ (1261.24) ions containing a palmitoylation at both C3 and C4. b and y ions of interest mentioned above are circled in red.

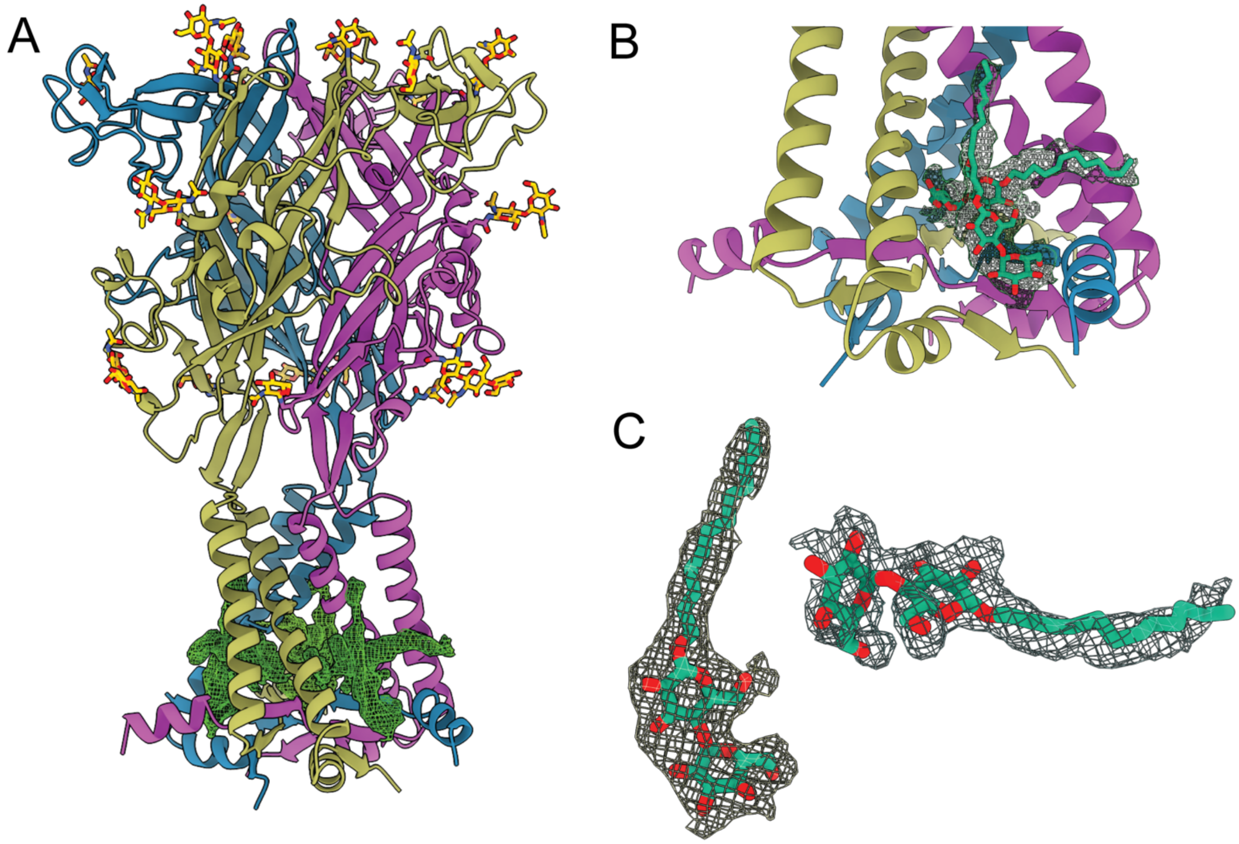


**Fig. S7. Cryo-EM density of the two DDM molecules bound in the cytoplasmic domain of $hP2X_4$ in the apo closed state. (A)** Ribbon representation of $hP2X_4$ in the apo closed state with the cryo-EM density for the DDM molecules shown in green mesh. The DDM molecules bind in the cytoplasmic fenestrations at the interface between neighboring protomers and interact with the cytoplasmic cap. **(B)** Magnified view of panel A, with both molecules of DDM now placed within their respective densities. **(C)** Magnified and isolated view of the two DDM molecules showing how well they fit within the density of the cryo-EM reconstruction.

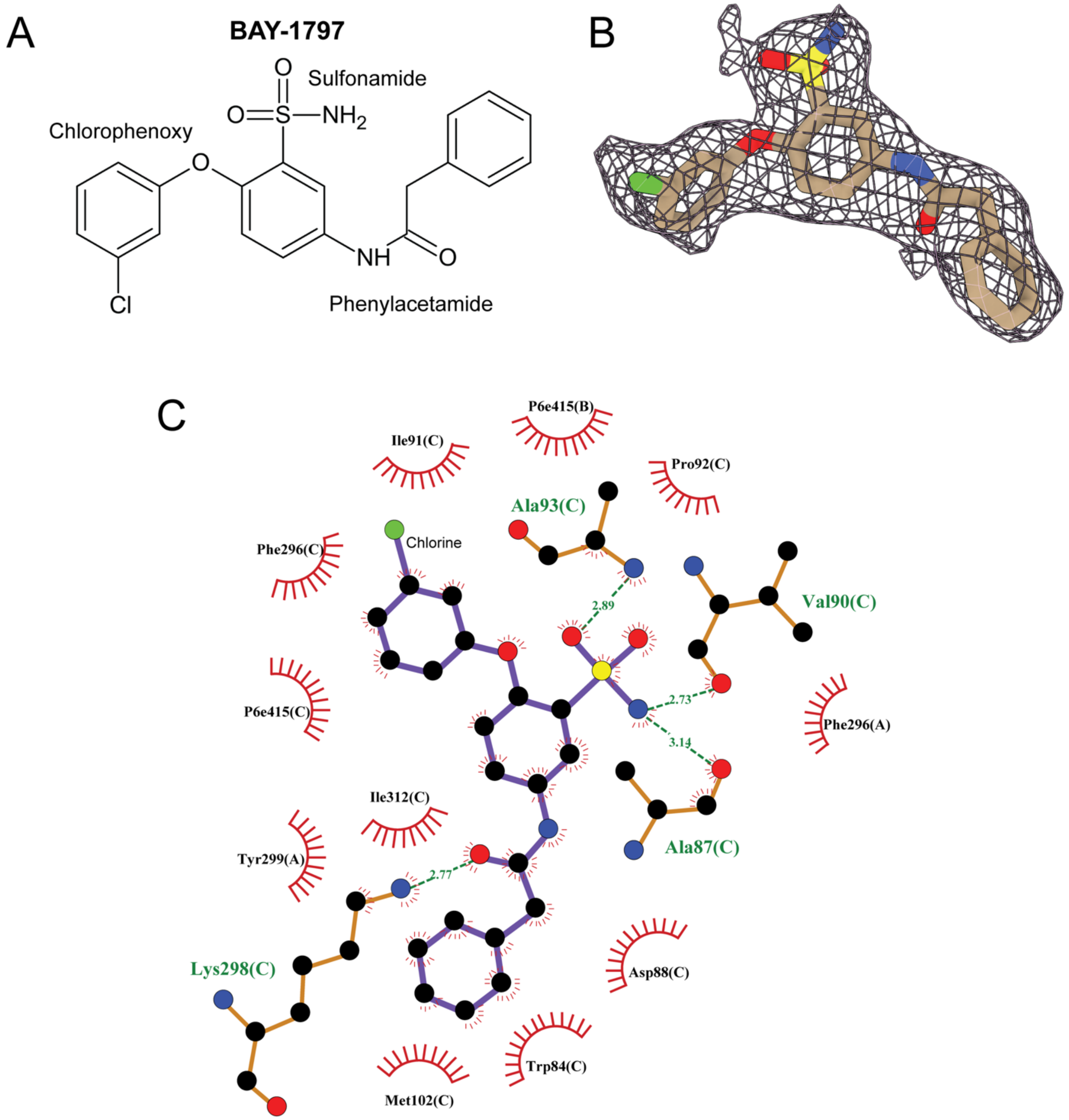


**Fig. S8. Chemical structure and functional group names of the $P2X_4$ allosteric antagonist, BAY-1797. (A)** Chemical structure of BAY-1797 highlighting the names of its functional groups. **(B)** Isolated view of BAY-1797 placed within the density of the cryo-EM reconstruction of the BAY-1797-bound inhibited state structure of $hP2X_4$. BAY-1797 is colored by atom: carbon atoms shown in tan, oxygen atoms in red, nitrogen atoms in blue, chlorine atom in green, and sulfur atom in yellow. **(C)** LigPlot diagram for BAY-1797 bound to $hP2X_4$ (55).

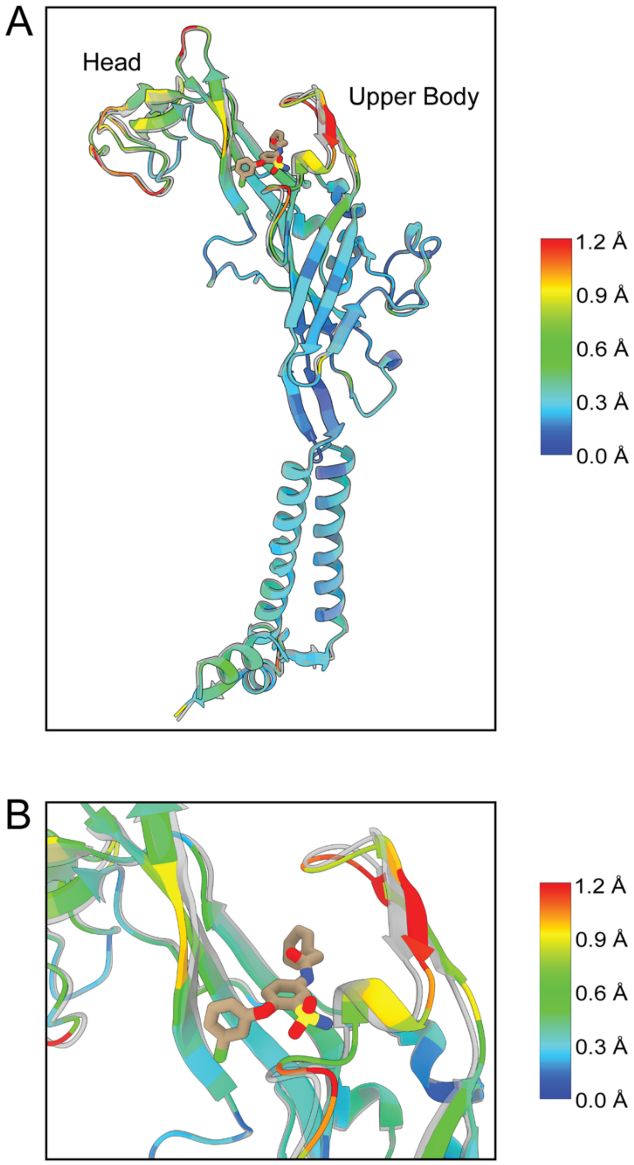


**Fig. S9. Alignment of hP2X4 in the apo closed state with hP2X4 in the BAY-1797-bound inhibited state. (A)** Structural alignment of a single protomer of hP2X4 in the apo closed state (colored by RMSD) with a single protomer of hP2X4 in the BAY-1797-bound inhibited state (transparent grey). The RMSD is color-coded to highlight the differences. The structures are very similar except for the expansion of the allosteric pocket of the binding site to accommodate BAY-1797. **(B)** Magnified view of panel A highlighting how the expansion of the allosteric pocket to accommodate BAY-1797 binding involves movements in the upper body domain. BAY-1797 molecules are colored by atom: carbon atoms shown in tan, oxygen atoms in red, nitrogen atoms in blue, chlorine atoms in green, and sulfur atoms in yellow.

**Table S1. Cryo-EM collection, refinement, and validation statistics.**

| | Apo closed hP2X4 (EMD-44799) (PDB: 9BQH) | ATP-bound hP2X4 (EMD-45177) (PDB: 9C48) | BAY-1797-bound hP2X4 (EMD-44800) (PDB: 9BQI) |
|---|---|---|---|
| **Data collection and processing** | | | |
| Magnification (kx) | 130 | 130 | 130 |
| Voltage (kV) | 300 | 300 | 300 |
| Electron exposure ($e^{-}$/Å$^2$) | 43 | 43 | 45 |
| Movie frames | 50 | 50 | 50 |
| Defocus range (μm) | -0.9 to -1.5 | -0.9 to -1.5 | -0.7 to -1.4 |
| Pixel size (Å) | 0.648 (0.324 super-res) | 0.648 (0.324 super-res) | 0.647 (Hardware Binned) |
| Symmetry imposed | C3 | C3 | C3 |
| Initial micrographs (no.) | 15,582 | 15,582 | 8,127 |
| Final micrographs used (no.) | 14,498 | 14,498 | 7,858 |
| Initial particle images (no.) | 2,225,406 | 2,225,406 | 1,475,108 |
| Final particle images (no.) | 627,118 | 122,405 | 239,914 |
| Map resolution (Å) | 2.27 | 2.40 | 2.55 |
| FSC threshold | (0.143) | (0.143) | (0.143) |
| Map resolution range (Å) | 1.4 - 25.9 | 1. 4 - 32.8 | 1.7 - 10 |
| **Refinement** | | | |
| Initial model used (PDB code) | 4DW0 | 5SVL | 4DW0 |
| Model resolution (Å) FSC threshold | 2.36 (0.143) | 2.37 (0.143) | 2.52 (0.143) |
| Map sharpening *B* factor (Å$^2$) | 64.3 | 75.3 | 82.8 |
| **Model composition** | | | |
| Non-hydrogen atoms | 9819 | 8481 | 9611 |
| Protein Residues | 1122 | 993 | 1122 |
| Ligands | 36 | 36 | 33 |
| Waters | 390 | 249 | 215 |
| *B* factors (Å$^2$) | | | |
| Protein | 10.89/66.71/29.29 | 37.5/138/64.0 | 38.92/80.83/47.78 |
| Ligand | 29.90/70.65/50.58 | 45.3/112/74.5 | 44.57/79.14/55.74 |
| Nucleotide | NA | NA | NA |
| Water | 9.80/39.95/21.72 | 37.2/73.5/50.2 | 39.55/65.83/47.43 |
| R.m.s. deviations | | | |
| Bond lengths (Å) | 0.005 (0) | 0.005 (0) | 0.004 (0) |
| Bond angles (°) | 0.625 (0) | 0.847 (0) | 0.601 (0) |
| Validation | | | |
| MolProbity score | 1.06 | 1.43 | 1.08 |
| Clash score | 2.28 | 5.35 | 2.89 |
| Poor rotamers (%) | 0 | 0 | 0 |
| Ramachandran plot | | | |
| Favored (%) | 97.76 | 97.26 | 98.3 |
| Allowed (%) | 2.24 | 2.74 | 1.70 |
| Disallowed (%) | 0 | 0 | 0 |
